# Adverse childhood experiences: associations with educational attainment and adolescent health, and the role of family and socioeconomic factors. Analysis of a prospective cohort study

**DOI:** 10.1101/612390

**Authors:** Lotte C Houtepen, Jon Heron, Matthew J Suderman, Abigail Fraser, Catherine R Chittleborough, Laura D Howe

**Author notes:** Corresponding author. MRC Integrative Epidemiology Unit, University of Bristol, Oakfield House, Bristol BS8 2BN, UK.

## Abstract

**Background:** Experiencing multiple adverse childhood experiences (ACE) is a risk factor for many adverse outcomes. However, the role of family and socioeconomic factors in these associations is often overlooked.

**Methods and findings:** Using data from the Avon Longitudinal Study of Parents and Children, we assess associations of ACE between birth and 16 years (sexual, physical or emotional abuse, emotional neglect, parental substance abuse, parental mental illness or suicide attempt, violence between parents, parental separation, bullying, and parental criminal conviction) with educational attainment at 16 years (n=9,959) and health at age 17 years (depression, obesity, harmful alcohol use, smoking and illicit drug use, n=4,917). We explore the extent to which associations are robust to adjustment for family and socioeconomic factors, whether associations differ according to socioeconomic factors, and estimate the proportion of adverse educational and health outcomes attributable to ACE, family or socioeconomic measures.

There were strong associations of ACE with lower educational attainment and higher risk of depression, drug use and smoking. Associations with educational attainment attenuated after adjustment but remained strong. Associations with depression, drug use and smoking were not altered by adjustment. Associations of ACE with harmful alcohol use and obesity were weak. We found no evidence that associations differed by socioeconomic factors. Between 5-15% of the cases of adverse educational and health outcomes occur amongst people experiencing 4+ ACE, and between 1-19% occur in people whose mothers have a low level of education.

**Conclusions:** This study demonstrates strong associations between ACE and lower educational attainment and worse health that are independent of family and socioeconomic factors. Our findings imply that interventions that focus solely on ACE or solely on socioeconomic deprivation, whilst beneficial, would miss most cases of adverse educational and health outcomes. Intervention strategies should therefore target a wide range of relevant factors, including ACE, socioeconomic deprivation, parental substance use and mental health.

## Introduction

There is increasing awareness of the role that adverse childhood experiences (ACE) can play in influencing educational attainment, physical and mental health.[1–6] ACE is rising rapidly on policy agendas and there is a drive within public health and education to prevent ACE, develop and implement interventions to improve resilience, and promote ACE-aware services.[7–10]

The definition of ACE varies between studies and is the subject of debate[11], but the adversities most commonly studied include child maltreatment (e.g. emotional, physical and sexual abuse, physical or emotional neglect) and measures of household dysfunction (e.g. violence between parents, parental separation and parental substance misuse, mental illness or criminal behaviour). It is well established that these adverse experiences are not randomly distributed across a population; socioeconomic disadvantage is a strong risk factor for ACE.[12–14] In the Avon Longitudinal Study of Parents and Children (ALSPAC), we have previously shown that young people from families with low social class have twice the prevalence of four or more ACE compared with young people from high social class families.[15] There is some debate about whether poverty should itself be considered an ACE; advocates point to the link between poverty and ACE and the advocacy advantages of having poverty included in the definition of ACE[16], whereas detractors view poverty as a structural issue, and highlight that ACE occur across the socioeconomic spectrum.[17] Here, we view socioeconomic disadvantage and ACE as separate phenomena.

Despite socioeconomic disadvantage being a major risk factor for ACE, the conversation about ACE rarely focuses on the inter-relationships between ACE and socioeconomic conditions or other family-level factors, and the implications of this for policy.[18] There is an extensive literature on factors that promote resilience to ACE[19–21], but there is relatively little focus in the literature on whether associations between ACE and adverse outcomes are weaker in children from socioeconomically advantaged families (one way of conceptualising resilience).

In this paper, we use data from a UK prospective cohort study to examine the associations of ACE from 0-16 years with educational attainment at 16 years and markers of adolescent health and health-related behaviours at age 17 years (depression, obesity, harmful alcohol use, smoking and illicit drug use). We assess the degree to which these associations are robust to adjustment for a wide range of family and socioeconomic characteristics, and we test the hypothesis that associations between ACE and education and health outcomes will be stronger in people from families with low socioeconomic position. We calculate population attributable fractions for each outcome for ACE and for several socioeconomic and family-related measures, to assess the relative contributions of each of these factors to adverse educational and health outcomes, with the motivation of understanding the proportion of cases of these adverse outcomes that could potentially be prevented by interventions focused solely on ACE.

## Methods

### Participants

The Avon Longitudinal Study of Parents and Children (ALSPAC) is a prospective, population-based birth cohort study that recruited 14,541 pregnant women resident in Avon, UK, with expected delivery dates between the 1st April 1991 and 31st December 1992.[22, 23] The mothers, their partners and the child have been followed-up using clinics, questionnaires and links to routine data. The study website contains details of all the data that is available through a fully searchable data dictionary: http://www.bris.ac.uk/alspac/researchers/data-access/data-dictionary/. Ethical approval for this study was obtained from the ALSPAC Law and Ethics Committee and the Local Research Ethics Committees.

To ensure sufficient data to inform multiple imputation, we excluded children with data on fewer than 10% of the ACE questions (n=2604). After this exclusion, 11,935 participants remained. We created separate analysis samples for analysis of educational attainment and health outcomes by restricting to participants with at least one outcome measure; we also excluded one child from within each twin pair (n=152) to maintain independence of observations. This resulted in final sample sizes of 9,959 for educational outcomes (obtained through linkage to routine data) and 4,917 for health outcomes (assessed at a research clinic).

### Adverse childhood experiences

Data on multiple forms of ACE were reported by both participants themselves and their mothers at multiple time points. Full details of the derivation of ACE measures has been described previously.[15] Briefly, dichotomous constructs indicating exposure to adversities between birth and 16 years were created for the ten ACE that are included in the World Health Organization ACE international questionnaire[24] (sexual abuse, physical abuse, emotional abuse, emotional neglect, parental substance abuse, parental mental illness or suicide attempt, violence between parents, parental separation, bullying and parental criminal conviction). The definitions are described in Supplemental Table 1. Most ACE data were collected prospectively, but some (in particular, sexual abuse) included retrospective reports.

### Educational attainment

ALSPAC data were linked to the National Pupil Database (NPD). This is a governmental database providing data on pupil level attainment in state funded schools in England. General Certificate of Secondary Education (GCSE) examinations are sat during the 11th year of compulsory schooling when children are aged 15/16 (years 2007–2009 for the ALSPAC cohort). Pupils study up to 12 subjects (8 on average). The subjects are graded individually on a scale of A* (highest) to G (lowest). For this analysis, we used a dichotomous indicator of less than five ‘good’ GCSEs (five or more grades A*-C including English and Mathematics), which is a widely-used benchmark of academic achievement in the UK and a requirement for entry into many further education courses.

### Health and health-related behaviours at age 17

During a research clinic at age 17 years (mean 17 years and 9 months, SD 4 months), weight was measured to the nearest 50g using the Tanita Body Fat Analyser (Model TBF 401A), with participants in underwear or light clothing and footwear removed. Height was measured using a Harpenden stadiometer to the last complete mm, with participants unshod. Body mass index was calculated as weight in kilograms divided by height in metres squared. Substance use and mental health were assessed using self-administered computer-assisted interviews. We derived the following dichotomous indicators:

- Obesity: Body mass index (BMI) was converted into sex- and age-specific Z-scores relative to UK 1990 population reference data. These Z-scores were used to define obesity based on published BMI Z-score cut-offs from the International Obesity Task Force (BMI-Z ⍰ 2.212 for boys and BMI-Z ⍰ 2.195 for girls).
- Regular smoking: We created a dichotomous indicator of smoking weekly or more versus no smoking or smoking less than weekly.
- Harmful drinking: ⍰ 16 on the 10-item alcohol use disorders identification test (audit).[25]
- Depression: based on the clinical interview schedule-revised (CIS-R), defined as meeting the depression diagnosis criteria of the international classification of diseases, 10th revision.[26]
- Illicit drug use: problematic cannabis use or, in the past 12 months, any use of any of the following substances: cocaine, amphetamines, inhalants, sedatives, hallucinogens, or opioids. Problematic cannabis use was measured using the six-item cannabis abuse screen test[27], which assesses cannabis consumption in the previous 12 months and focuses on difficulties controlling use and associated health and social impairment. All items are answered on a 5-point scale (0 never, 1 rarely, 2 from time to time, 3 fairly often, and 4 very often). A response of fairly often or very often to any of the six items was used to indicate problem cannabis use.

### Confounders

At enrolment and prior to delivery, several self-report questionnaires were administered that measured socio-economic, family and (mental) health variables. Based on these parental questionnaires, the following covariables were included in the analysis: mother’s home ownership status during pregnancy (Mortgaged/ Owned/ Council rented/ Furnished private rental/ Unfurnished private rental/ Housing authority rented/ Other), mother and partner’s highest educational qualification (CSE/ Vocational/ O-level/ A-level/ Degree), household social class (highest of mother and partner social class according to the Registrar General’s Social Classes: professional/ managerial and technical/ skilled non-manual/ partly skilled/ unskilled), parity, maternal report of child’s ethnicity (white/non-white), mother’s age at delivery (in years), mother’s marital status during pregnancy (Never married/ Widowed/ Divorced/ Separated/ 1^st^ marriage/ Marriage 2 or 3), mother’s depression score (EPDS) at 18 and 32 weeks gestation and mother’s partner depression score (EPDS) at 18 weeks gestation. More details on these variables and their distributions are available in Supplemental Table 2.

### Missing data

Due to the derivation of ACE measures from multiple questionnaires and clinics over a long time period (birth-23 years), no participants had data on all of the individual questionnaire items, necessitating the use of multivariate multiple imputation. Ideally, we would impute missing values of each questionnaire item, but the lack of complete cases in combination with the high number of variables (>500 separate questions relating to ACE) led to convergence errors. Therefore, we adopted a pragmatic approach to imputation, adapted from the scale level imputation method proposed by Enders.[28] We derived a dichotomous construct indicating presence or absence of each ACE. If a participant responded to 50% or more of the questions related to a given ACE, we used these data to create the dichotomous indicator. If the participant responded to less than 50% of the questions, we set the dichotomous indicator to missing. We derived a cumulative adversity measure (ACE-score) by summing exposure to the ten classic ACE, defining four categories (0, 1, 2-3 and more than 4 ACE). Due to the considerably larger sample size available for educational attainment than for health measures, these groups of outcomes were considered in separate imputation models. As there are some sex differences in ACE prevalence (Supplemental Table 2) and potentially higher order interactions between sex and adversity that we want to preserve, males (education n=5,023, health n=2,163) and females (education n=4,936, health n=2,754) were imputed separately before appending the two datasets before analysis. The dichotomous ACE indicators and the ACE score were included in multiple imputation models, along with outcome variables and auxiliary variables likely to predict either missingness, ACE exposure or health status (sociodemographic indicators, adversity measures from before the child’s birth, and additional education and health variables – additional details in Supplementary Table 3 and full details in previous publication[15]). For both males and females, 90 imputed datasets were created using the mice package in R3.3.1 with 30 iterations per dataset. For secondary analyses exploring interactions between ACE and parental social class or maternal education, imputation models were re-run stratified by dichotomous indictors of i) parental social class (manual versus non-manual, highest social class of mother or partner) and ii) maternal education (CSE, vocational education or lower versus O-level, A-level, degree or higher).

### Statistical analyses

All statistical modelling was done in R version 3.3.1 unless otherwise specified, using binary logistic regression models for all outcomes. Associations of each separate ACE and the ACE-score with each outcome were assessed in a basic model (adjusted for sex) as well as a fully-adjusted model (adjusted for home ownership, maternal and partner education, household social class, parity, ethnicity, maternal age, maternal marital status, maternal and partner depression during pregnancy). For the imputed data, the logistic regression results were obtained by averaging across the results from each of the 90 imputed datasets using Rubin’s rules. This procedure appropriately modifies the standard errors for regression coefficients (used to calculate p-values and 95% confidence intervals) to take account of uncertainty in both the imputations and the estimate. Likelihood ratio test statistics were combined using an approximation proposed by Meng and Rubin.[29] As a sensitivity analysis, we replicated these analyses in people with ‘complete’ data, i.e. participants who responded to more than 50% of the questionnaire items for all ACE and who had data on the outcomes.

To examine whether the associations differed according to sex we used likelihood ratio tests for interaction, and if applicable report the results of sex stratified analyses. We also used likelihood ratio tests to assess interactions between ACE and manual versus non-manual parental social class and low versus high maternal education.

Prevalence of the outcomes across exposure categories and risk differences and ratios were estimated in the imputed data using the ‘mim: glm’ command in Stata version 15. Population attributable fractions (PAF) were estimated using the formula 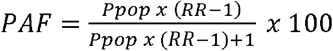 where *Ppop* is the proportion of exposed participants and *RR* is the risk ratio. PAF estimates the percentage of cases of the outcome that would be prevented if the exposure was eliminated (assuming causality and absence of bias). Alternatively, it could be conceptualized as the percentage of people who go on to develop the adverse outcome that would be included in interventions targeted at the risk factor of interest. This analysis was performed for each of our binary outcomes for the following exposures, which were selected to represent a range of potential ways of identifying high risk groups, most of which have a prevalence broadly similar to 4 or more ACE: 4 or more ACE (19%), low maternal education (CSE, vocational qualifications, or lower; 30%), manual social class (classes IIIm, IV or V in the 1991 UK Office of Population Censuses and Surveys classification; 23%), maternal depression in pregnancy (score of 12 or more on the EPDS on either of the two pregnancy questionnaires; 21%), any self-reported maternal smoking during pregnancy (26%), social housing during pregnancy (15%) and maternal age less than 20 years (4%).

### Data availability

ALSPAC data are available by application to the study executive committee, for details please see http://www.bristol.ac.uk/alspac/researchers/access/. Full details of the derived measures of ACE used in this paper are published in a Data Note[15], and these data, including analysis scripts for multiple imputation, are available on request from the ALSPAC study data team.

## Results

84% of ALSPAC participants were exposed to at least one ACE; 23.6% were exposed to one ACE, 36.5% to two or three ACE, and 23.8% to four or more ACE (Table 1). The distribution of the ACE score was similar in males and females (Supplementary table 2). The prevalence of individual ACE ranged from 4.1% for sexual abuse to 48.6% for parental mental health problems. The prevalence of most individual ACE was similar in males and females, apart from sexual abuse, which was reported by 2.3% of males and 6.0% of females (Supplementary table 2). Consistent with a higher rate of missing data in more deprived participants who are more likely to drop out from the cohort[30], the ACE prevalence estimates were higher in the imputed data compared with the raw data[15] (Supplemental Table 4).

Just over half (54.5%) of participants received five or more good GCSEs (Table 1). The prevalence of health outcomes at age 17 years was 7.3% for obesity, 8.7% for depression, 19.5% for smoking, 16.1% for drug use, and 10.9% for harmful alcohol use (Table 1).

**Table 1.** Participant characteristics. Characteristics of the participants included in analyses, using data from multivariate multiple imputation. N= 9,959 apart from for health outcomes, where N=4,917

|  | Mean (SE) for continuous variables<br>% for categorical variables |
| --- | --- |
| ACE-score: 0 | 16.1 |
| 1 | 23.6 |
| 2 to 3 | 36.5 |
| 4+ | 23.8 |
| Physical abuse | 19.0 |
| Sexual abuse | 4.1 |
| Emotional abuse | 23.9 |
| Emotional neglect | 23.9 |
| Bullying | 26.2 |
| Violence between parents | 25.3 |
| Parental substance abuse | 15.1 |
| Parental mental health problems or suicide attempt | 48.6 |
| Parental criminal conviction | 10.5 |
| Parental separation | 33.8 |
| 5+ GCSEs including math and English at grades A*-C | 54.5 |
| Obesity at age 17<sup>†</sup> | 7.3 |
| Depression at age 17<sup>†</sup> | 8.7 |
| Smoking at age 17<sup>†</sup> | 19.5 |
| Illicit drug use at age 17<sup>†</sup> | 16.1 |
| Harmful alcohol use at age 17<sup>†</sup> | 10.9 |

5+ GCSEs indicates five or more grades A\*-C including English and Mathematics from GCSE or equivalent examinations. Obesity is defined as BMI Z-score above cut-offs from the International Obesity Task Force (BMI-Z ≥2.212 for boys and BMI-Z ≥ 2.195 for girls). Depression is defined as meeting the depression diagnosis criteria of the international classification of diseases, 10th revision on the clinical interview schedule-revised (CIS-R). Smoking is defined as weekly smoking. Illicit drug use is defined as problematic cannabis use

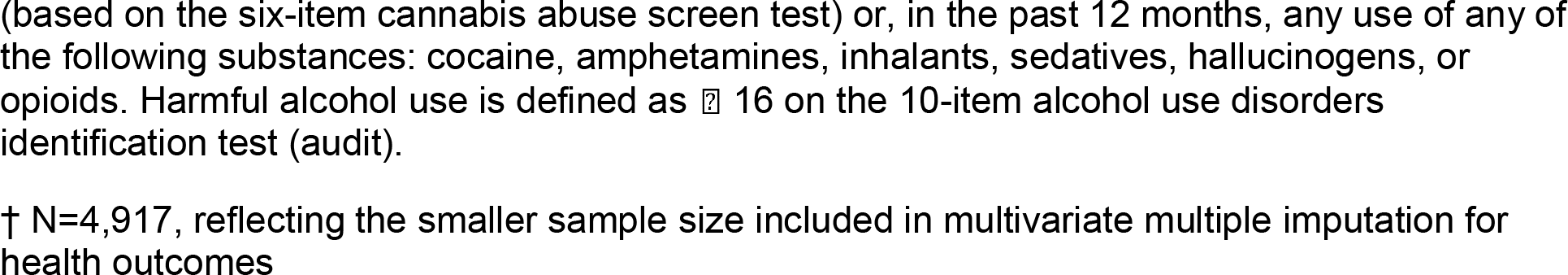

There was no evidence for sex interactions in most of our analyses (Supplemental Table 5); therefore, the results of the sex stratified analyses are only mentioned when there was evidence for a sex interaction (p<0.05 on likelihood ratio tests for interaction). We describe the results from analysis of multiply imputed data as our main results; results from complete case analysis were generally closer to the null than in imputed data, but the overall picture of results was similar (Supplementary Table 5).

### Association between ACE and educational attainment

Figure 1 and Supplementary Table 5 show the results for the associations of the ACE score and each individual ACE with educational attainment. The ACE score was strongly associated with lower educational attainment. Associations were apparent even when comparing those experiencing only one ACE to those with no ACE (e.g. OR for less than five good GCSEs 1.37, 95% CI: 1.15 to 1.63 after adjustment for confounders) and were stronger for each increasing value of the ACE score. Experiencing four or more ACE was associated with 50% higher odds of obtaining less than five good GCSEs (OR 1.46, 95% CI: 1.04 to 2.05) after adjustment for confounders (Supplemental Table 5).

Most of the individual ACE were associated with lower educational attainment. The strongest associations were seen for emotional neglect, e.g. (OR for less than five good GCSEs 1.81, 95% CI 1.49 to 2.2). Associations were weak or absent for physical abuse. For all ACE and all measures of educational attainment, associations were considerably weaker after adjustment for confounders. For bullying, there was evidence of a sex interaction, with the associations with less than five good GCSEs being stronger in females compared with males (Supplemental Table 5).

**Figure 1.**
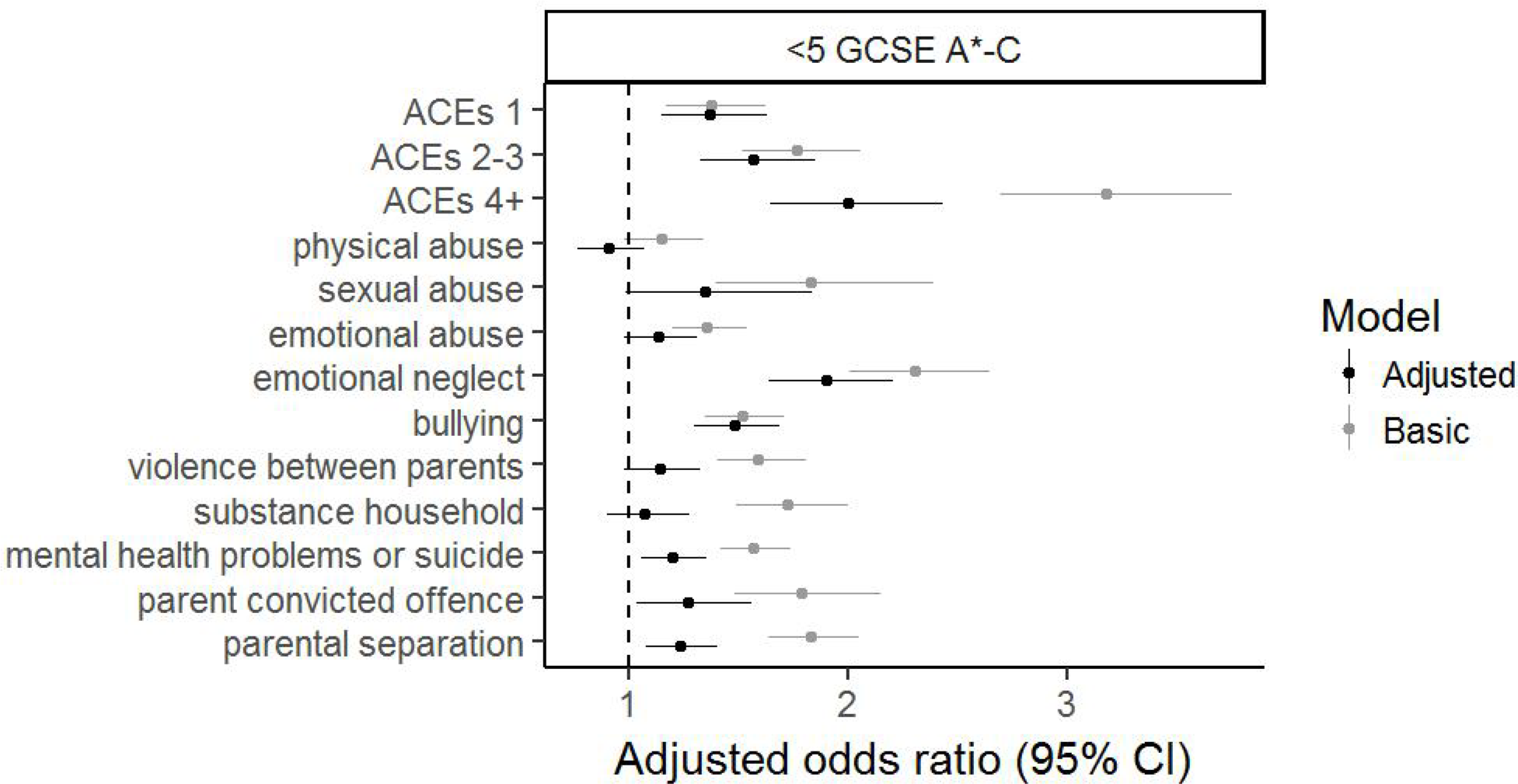
Forest plot for the associations of the ACE score and separate ACE with obtaining less than five GCSEs at A*-C, including maths and English. The reference category for each category of the ACE score (1, 2-3 and 4+) is experiencing 0 ACE. The basic model is adjusted for sex. The adjusted model additionally includes home ownership, maternal and partner education, household social class, parity, ethnicity, maternal age, maternal marital status, maternal and partner depression during pregnancy.

There was no consistent evidence of an interaction between ACE and parental social class or maternal education (Supplementary Tables 6 and 7; Supplementary Figures 1 and 2) in their relationship with educational outcomes.

### Association between ACE and health/health-related behaviours

The ACE score was strongly associated with depression, illicit drug use, and smoking (Figure 2, Supplementary Table 5). The group of participants who experienced one ACE had higher odds of all of these outcomes compared to those who experienced no ACE, but the confidence intervals included the null apart from for illicit drug use (OR after adjustment for confounders 1.4, 95% CI 1.0 to 2.0). People who experienced two to three, or four or more ACE were more likely to be depressed, use illicit drugs, and smoke, with associations generally stronger in the four or more ACE group. People who experienced 4+ ACE were more than twice as likely to smoke (OR 2.3, 95% CI 1.7 to 3.2), to be depressed (OR 2.5, 95% CI 1.6 to 3.0), and three times more likely to use illicit drugs (OR 3.0, 95% CI 2.1 to 4.2) compared to people who experienced no ACE (Supplementary Table 5).

Obesity and harmful alcohol consumption demonstrated weak associations with the ACE score; OR for 4+ ACE compared with no ACE was 1.4 for obesity (95% CI 0.9 to 2.2) and 1.4 for harmful alcohol consumption (95% CI 0.9 to 2.0).

Examining individual ACE revealed strong associations of: i) physical abuse with depression, illicit drug use, and smoking, ii) sexual abuse with depression and smoking, iii) emotional abuse with illicit drug use and smoking, iv) bullying with depression, v) violence between parents with illicit drug use and smoking, vi) parental substance abuse with harmful alcohol use, illicit drug use and smoking, vii) parental mental health problems with depression, illicit drug use and smoking, vii) parental separation with illicit drug use and smoking. Thus overall, the patterns mirrored those of the ACE score – associations were strong for depression, illicit drug use and smoking, and weak or absent for obesity and harmful alcohol use. The strongest associations tended to be seen for physical and sexual abuse, although very strong associations were seen for parental substance use in relation to substance use outcomes in the offspring.

**Figure 2.**
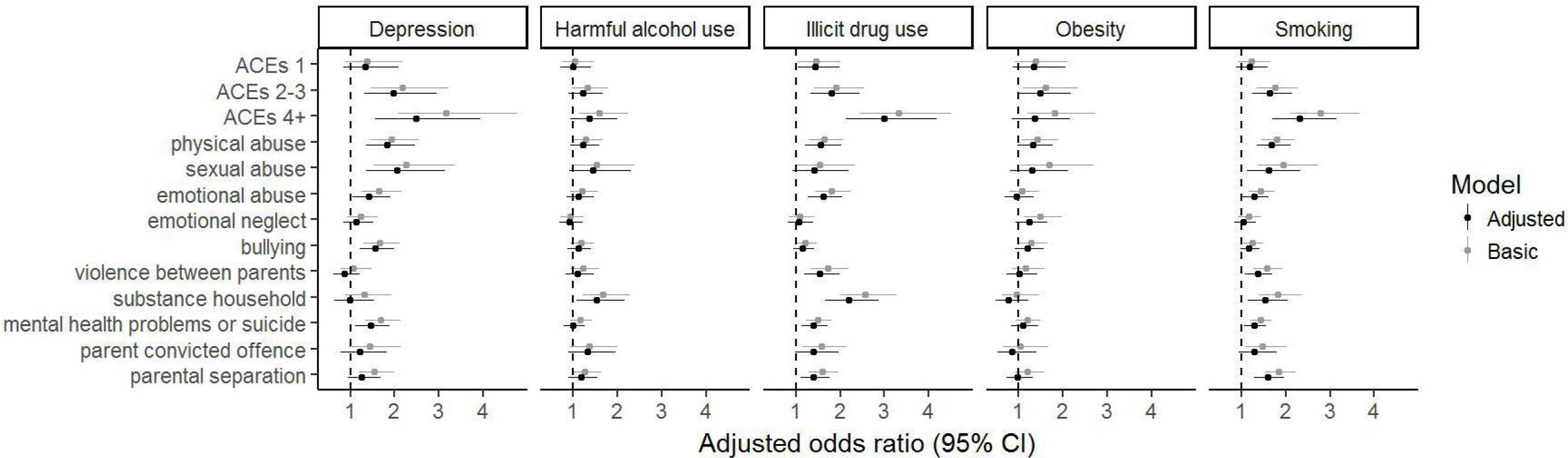
Forest plots for the associations of the ACE score and separate ACE with poor health outcomes. The reference category for each category of the ACE score (1, 2-3 and 4+) is experiencing 0 ACE. The basic model is adjusted for sex. The adjusted model additionally includes home ownership, maternal and partner education, household social class, parity, ethnicity, maternal age, maternal marital status, maternal and partner depression during pregnancy.

In contrast with the educational outcomes, adjustment for sociodemographic confounders did not markedly attenuate the associations between ACE and health/behavioural outcomes, and in some instances, adjustment resulted in stronger associations.

The only sex interaction was for exposure to parental substance abuse and illicit drug use (Supplemental Table 5), with stratified analyses indicating a stronger association in females than males (females OR=2.8 95% CI 1.9 to 4.1; males OR=1.7 95% CI 1.1 to 2.6).

Consistent with the educational outcomes, there was no evidence that associations between ACE and health and behavioural outcomes differed according to parental social class or maternal education (Supplementary Tables 6 and 7 and Supplementary Figures 1 and 2).

### Population attributable fractions

Differences in risk of achieving less than five good GCSEs (Table 2) ranged from 11% (95% CI 9% to 14%) for maternal depression during pregnancy to 35% (95% CI 32% to 37%) for social housing. All but one sociodemographic factor (maternal depression) had risk differences (RDs) that were higher than for 4+ ACE (RD 18%, 95% CI 15% to 20%). The lowest PAF was for maternal age less than 20 years (2%), reflecting the low prevalence of this risk factor. The highest PAF was for maternal education (19%). This compares to 9% for 4+ ACE.

**Table 2.** Associations of ACE and various sociodemographic markers with educational attainment on the risk difference scale, and population attributable fractions. PAF = population attributable fraction; the proportion of the people experiencing <5 good GCSEs who also experienced the ‘exposure’ (ACE or sociodemographic variable); can be interpreted as the proportion of the cases of <5 GCSEs that could be prevented if the exposure was eliminated, assuming causality. Note that the reference categories in this table differ from those in other parts of the manuscript; here, the reference category is all other participants apart from those with the exposure, for example the reference category for 4+ ACE here is <4 ACE.

|  | *Prevalence of exposure* | *Less than 5 good GCSEs* |  |  |  |
| --- | --- | --- | --- | --- | --- |
|  |  | Prevalence in exposed | Prevalence in unexposed | Risk difference (95% CI) | PAF |
| 4 or more ACEs | 19.0% | 59.1% | 41.2% | 18%<br>(15% to 20%) | 9% |
| Low maternal education (CSE, vocational, or lower) | 29.6% | 65.6% | 36.8% | 29%<br>(27% to 31%) | 19% |
| Manual social class (III manual and lower) | 22.9% | 64.8% | 39.4% | 25%<br>(23% to 28%) | 13% |
| Maternal depression in pregnancy | 20.5% | 54.3% | 43.1% | 11%<br>(9% to 14%) | 5% |
| Any maternal smoking in pregnancy | 25.7% | 59.6% | 40.4% | 19%<br>(17% to 21%) | 11% |
| Social housing | 14.5% | 75.3% | 40.4% | 35%<br>(32% to 37%) | 11% |
| Mother aged 19 years or lower at birth | 3.6% | 74.6% | 44.5% | 30%<br>(25% to 35%) | 2% |

The highest obesity PAFs were seen for socioeconomic markers; e.g. the PAFs for social housing and low maternal education were 14% and 13% respectively, compared with 5% for 4+ ACE (Table 3). In contrast, PAFs for illicit drug use and depression were highest for 4+ ACE, with high PAFs also seen for maternal smoking in pregnancy and relatively low PAFs for socioeconomic and other family measures. For example, the PAFs of depression for 4+ ACE, maternal smoking during pregnancy and manual social class were 14%, 10% and 3% respectively.

Harmful alcohol use and smoking exhibited a different pattern again; the highest PAF for each of these outcomes was for maternal smoking during pregnancy – 11% for harmful alcohol use and 15% for smoking. PAFs for 4+ ACE were 6% for harmful alcohol use and 12% for smoking. Socioeconomic markers had lower PAFs for harmful alcohol use, but PAFs for smoking were similar to the PAF for 4+ ACE, e.g. PAF for manual social class in relation to smoking was 10%.

**Table 3.**
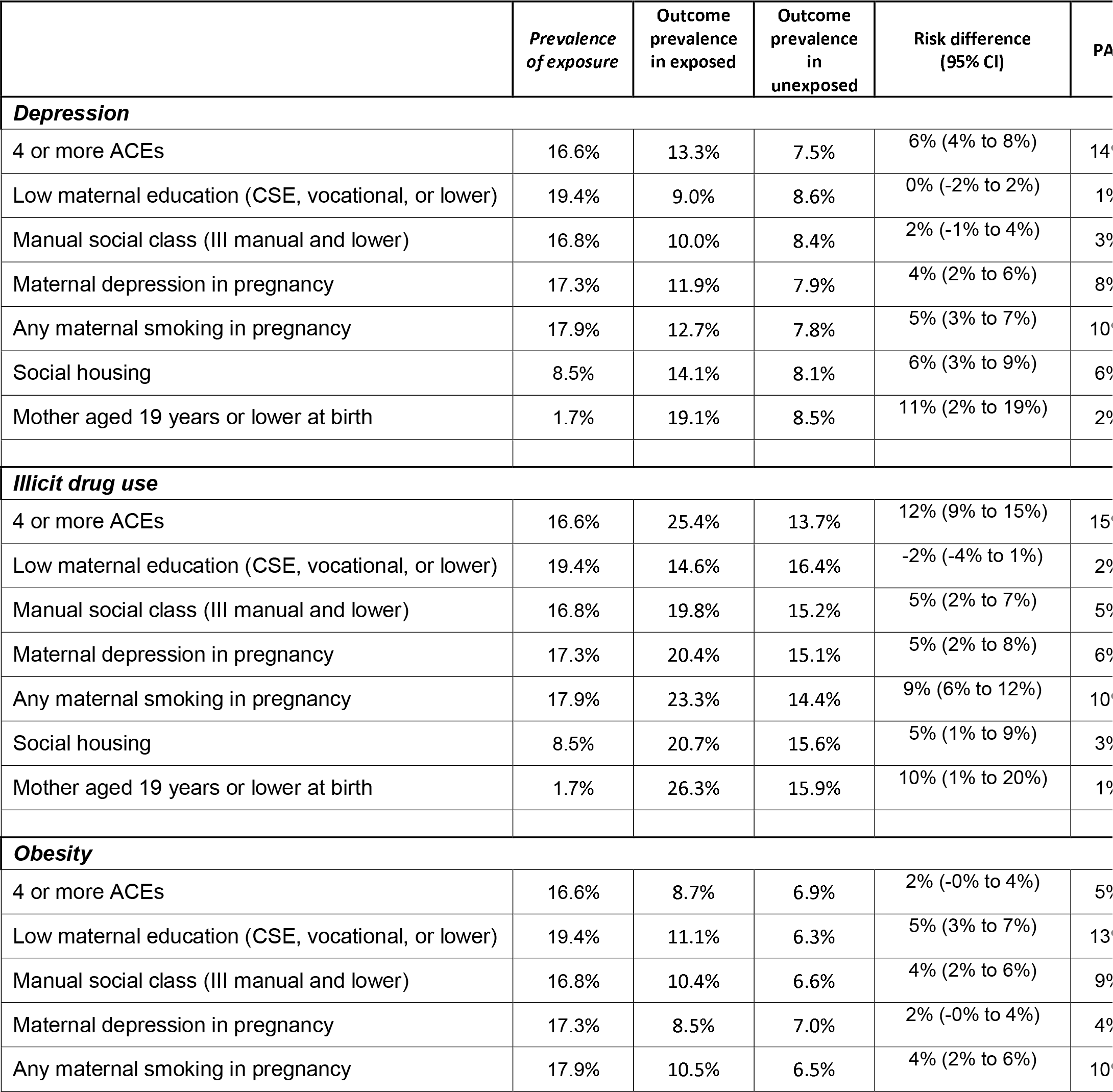
Associations of ACE and various sociodemographic markers with health and health risk behaviours on the risk difference scale, and population attributable fractions. PAF = population attributable fraction; the proportion of the people experiencing each outcome (depression, illicit drug use, obesity, harmful alcohol use, or smoking) who also experienced the ‘exposure’ (ACE or sociodemographic variable); can be interpreted as the proportion of the outcome cases that could be prevented if the exposure was eliminated, assuming causality. Note that the reference categories in this table differ from those in other parts of the manuscript; here, the reference category is all other participants apart from those with the exposure, for example the reference category for 4+ ACE here is <4 ACE.

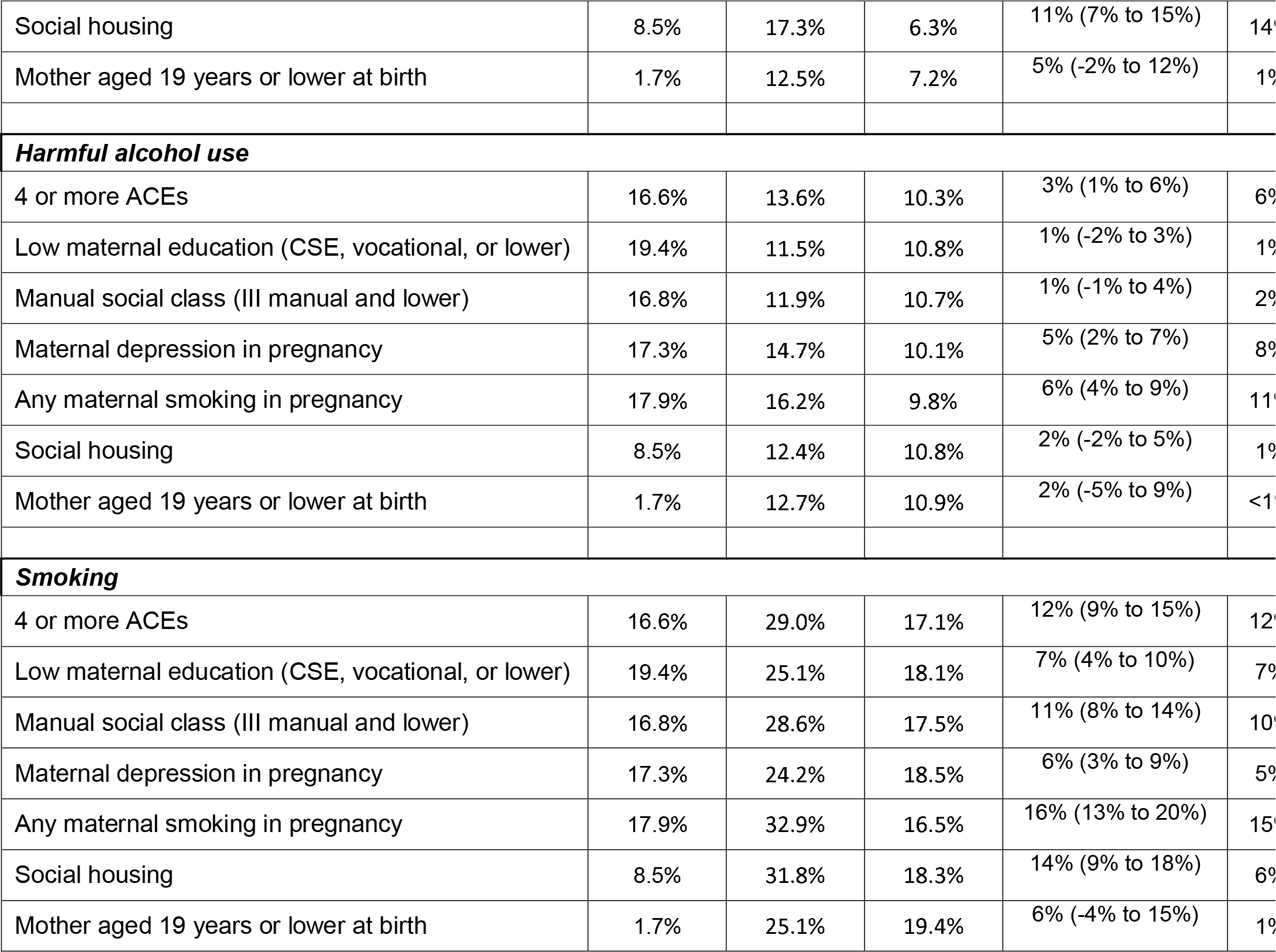

## Discussion

In this UK cohort, where 84% of participants were exposed to at least one ACE and 24% were exposed to four or more ACE, we find evidence that ACE – both when considered together as an ACE score and separately as individual ACE – are associated with lower educational attainment and worse health and health-related behaviours. Adjustment for a wide range of family and socioeconomic variables reduced the magnitude of associations of ACE with educational attainment by approximately half. However, adjusted associations between ACE and educational attainment were still strong, and similar in magnitude to the strongest associations between ACE and health/health-related behaviours. Adjustment for confounders did not attenuate associations between ACE and health and health-related behaviours. We found no evidence that higher SEP acted as a buffer to the adverse effects of ACE; associations between ACE and both educational and health outcomes were similar in adolescents with parents from manual and non-manual occupational social classes and for adolescents with low and high levels of maternal education. When calculating the proportion of cases of each outcome attributable to 4+ ACE or various family and socioeconomic measures, we found a different pattern of results across the outcomes. For education and obesity, the highest PAF were observed for socioeconomic markers. In contrast, PAFs for illicit drug use and depression were highest for 4+ ACE, and for harmful alcohol use and smoking the highest PAF was for maternal smoking during pregnancy.

The attenuation of ACE effects on education by half when adjusting for family and socioeconomic factors suggests that the family and socioeconomic context is responsible for a considerable proportion of these associations. However, most studies of the social and health sequalae of ACE do not include the broad range of confounders that we included. Consequently, they may be overestimating the impact of ACE. Although we adjusted for a wide array of factors, we are unlikely to have captured all relevant concepts, and our measurements will not perfectly capture the concepts of interest. For example, current housing tenure may not completely capture life course trajectories of housing tenure, and does not fully capture crowding, damp, residential instability, and other important aspects of housing. Therefore, residual confounding is likely to be present, and our estimates of the educational impact of ACE are likely to be overestimates. In contrast, our results indicate that, at least in this population, the associations of ACE with health and health-related behaviours in adolescence were not strongly affected by adjustment for sociodemographic confounders, suggesting that ACE are associated with these outcomes regardless of the family and sociodemographic setting in which they are experienced. The degree to which previous studies have adjusted for potential confounding variables varies considerably, with some studies making no attempt at adjustment.[6] Studies that do adjust for a range of factors differ in whether this adjustment leaves associations largely unchanged[31] or results in considerable attenuation.[1]

Our *a priori* hypothesis was that associations between ACE and adverse educational and health outcomes would be weaker in adolescents from high SEP families, as other aspects of a high SEP environment could act to mitigate the effects of ACE. We found no evidence to support this hypothesis; associations were of similar magnitude in manual and non-manual social class families and in families with high and low levels of maternal education. Other studies have also found no differences in associations between ACE and outcomes according to socioeconomic position[32] or race[31], and one study found either no difference in ACE-outcome associations according to income, or stronger associations in high-income groups.[14] Together, these findings support universal ACE prevention or support interventions, rather than focusing ACE initiatives only in low socioeconomic population groups.

Population attributable fractions (PAFs) estimate the proportion of an outcome that could be eliminated if the exposure is removed from the population. This interpretation of a PAF requires unrealistically strong assumptions about causality and lack of bias. Since the exposures we examine are inter-related, the PAFs should also not be considered in isolation, because a joint PAF for all exposures considered together is likely to be considerably smaller than implied by the individual PAFs. Nonetheless, the comparative magnitude of PAFs across exposures may be informative. Our results imply that ACE-focused interventions may have less impact on population-level educational attainment compared with interventions or policies that address socioeconomic disadvantage, whereas ACE-focused initiatives may yield the greatest population-level effects on depression and drug use in adolescents. The findings for smoking and harmful alcohol use may reflect family-level propensity for risky behavior and intergenerational transmission of behaviours. For all outcomes, PAFs were relatively low – ranging from 1 to 15% across all exposures, and 5 to 15% for four or more ACE. Thus, between 5-15% of the cases of low educational attainment and poor health/health-related behaviour occurred in the participants who experienced four or more ACE, meaning that interventions targeting subgroups based on solely exposure to ACE will fail to prevent most cases.

Within the ALSPAC cohort, there is a vast amount of information, mainly prospectively collected from both young people and their mothers, from various time points and life stages. This vast array of data, considered together, resulted in a higher prevalence of many ACE than is seen in some other studies. For example, in the Welsh ACE study, one in seven participants reported four or more ACE[33], compared with one in four in ALSPAC. Studies using a single retrospective questionnaire may be underestimating the prevalence of ACE. However, it is also possible that our cohort is identifying a set of people for whom experience of ACE has been less severe or of shorter duration than those identified in cross-sectional studies collecting retrospective reports of ACE exposure at a single time point in adults. There is evidence that retrospective and prospective reports of ACE capture largely non-overlapping groups of individuals, but that both groups are at risk from adverse outcomes.[34, 35]

Although using data from multiple questionnaires across a long period of time enabled us to capture a detailed picture of the cohort members’ experience of ACE, data missingness became a challenge. We assumed that the data are missing-at-random given the variables included in the imputation model. Although this assumption is untestable, it allows for maximum use of the available data, and we included a number of key sociodemographic variables in the imputation model to make this assumption more plausible. In general, we would anticipate that deviations from the missing at random assumption would lead to underestimation of the ACE prevalence and potentially bias associations of ACE with adverse outcomes towards the null.[30] ALSPAC is a geographical cohort based in the South West of England, and has slightly higher levels of socioeconomic advantage and lower levels of ethnic diversity than the national average, and this may affect the generalisability of our findings.

We used two ways of conceptualizing ACE – first as a score of the number of ACE experienced, and second considering each ACE separately. Although the ACE score is widely used and has advantages including the recognition that ACE tend to co-occur and the worst outcomes tend to be for people exposed to multiple ACE, it is also problematic in several ways. For example, it assumes that each ACE has the same magnitude and direction of association with the outcome[36]; our analysis of individual ACE demonstrates this not to be the case. Nonetheless, we opted to use the ACE score approach rather than alternative grouping methods such as factor analysis or latent class models[37], to be consistent with other studies. When analyzing each individual ACE, we did not adjust for other ACE as covariates. The rationale for this is that the causal structure linking multiple ACE is complex and largely unknown. Some adjustments would therefore be over-adjustment, removing some of the effect of interest.

Our results suggest strong associations of ACE with lower educational attainment and worse health and health-related behaviours in late adolescence that are robust to adjustment for a wide range of variables describing the family and socioeconomic context. However, our data indicate that people experiencing 4+ ACE contribute between 5-15% of the cases of adverse educational and health outcomes considered in this study. This implies that prevention of ACE or improved support for people who experience ACE, whilst beneficial, would not affect the vast majority of people experiencing adverse educational and health outcomes in adolescence. The loss of human potential associated with ACE has led to urgent calls for ACE-awareness and action, to ensure that young people reach their developmental potential. There has been an upsurge of discussion about ACE in both the research and policy spheres. Our results suggest that, while welcome, interventions targeted at ACE prevention/support should be considered alongside other risk factors, including socioeconomic deprivation, parental substance use and mental health.

## Acknowledgments

We are extremely grateful to all the families who took part in this study, the midwives for their help in recruiting them, and the whole ALSPAC team, which includes interviewers, computer and laboratory technicians, clerical workers, research scientists, volunteers, managers, receptionists and nurses.

