## Supplementary Material for "Adverse childhood experiences: associations with educational attainment and adolescent health, and the role of family and socioeconomic factors. Analysis of a prospective cohort study"

**Supplementary Information**

***Adverse childhood experiences (ACE) definitions***

The cut-offs for each individual question are described in (1). In short, the adverse childhood experiences were defined as:

1. adult in family was ever physically cruel towards or hurt the child (*physical abuse*);
2. ever sexually abused, forced to perform sexual acts or touch someone in a sexual way (*sexual abuse*);
3. parent was ever emotional cruel towards child or often said hurtful/insulting things to the child (*emotional abuse*);
4. child always felt excluded, misunderstood or never important to family, parents never asked or never listened when child talked about their free time (*emotional neglect*);
5. child was a victim of bullying on a weekly basis (*bullying*);
6. parents were ever affected by physically cruel behaviour by partner, or, ever violent towards each other, including hitting, choking, strangling, beating, shoving (*violence between parents*);
7. parent was a daily cannabis or any hard drug user, or, had an alcohol problem (*substance use in household*);
8. parent was ever diagnosed with schizophrenia or hospitalised for a psychiatric problem, or, during the first 16 years of the child’s life, parent had an eating disorder (bulimia or anorexia), used medication for depression or anxiety, attempted to commit suicide or scored above previously established cut-offs for depression (Edinburgh Postnatal Depression Scale (EPDS) >12 (2)) (*mental health problems or suicide*);
9. parent was convicted of an offence (*parent convicted*);
10. parents separated or divorced (*parental separation*);

*Supplemental Table 1 Phrasing and cut-off criteria for the ALSPAC adversity questions used to derive the ACE constructs used in this study*

| **Adverse childhood experience (ACE)** | **Phrasing** | **Criterion** | **Retrospective** |
| --- | --- | --- | --- |
| Physical abuse | Partner/respondent was physically cruel to child | yes | no |
|  | Adult in family pushed, grabbed, shoved/smacked to discipline respondent, before age of 11 | often | yes (asked at 22yrs) |
|  | When growing up people in respondent's family hit them so hard that it left them with bruises or marks | yes | yes (asked at 23yrs) |
|  | Adult in family kicked, punched, hit respondent (so hard it left bruises or marks), before age of 11 | yes | yes (asked at 22yrs) |
| Sexual abuse | Sexually abused | yes | no |
|  | When growing up someone molested respondent (sexually) | yes | yes (asked at 23yrs) |
|  | Touched in a sexual way by adult or older child, or was forced to touch adult or older child in a sexual way, before age of 11 | yes | yes (asked at 22yrs) |
|  | Adult or older child forced, or attempted to force, respondent into any sexual activity by threatening or holding respondent down or hurting respondent in some way, before age of 11 | yes | yes (asked at 22yrs) |
| Emotional abuse | Partner/respondent was emotionally cruel to child | yes | no |
|  | Adult in family shouted/ said hurtful or insulting things to respondent, before age of 11 | Very often | yes (asked at 22yrs) |
| Emotional neglect | Carer knows who friends are | never | no |
|  | Carer asks/starts conversation about free time/ what happened at school | never | no |
|  | Carer takes time to listen when teenager talks about what happened in free time | never | no |
|  | Discuss problems with anyone in their family | very difficult | no |
|  | Parent/carer talked about child's experiences at school/ friends/ things that are troubling | never | no |
|  | Child feels left out of things | always | no |
|  | Understood by parents | not | no |
|  | When growing up there was someone to take respondent to the doctor if needed | never | yes (asked at 23yrs) |
|  | Someone in family made child feel important or special, before 11 | never | yes (asked at 22yrs) |
|  | Carer knows what child does with other children | nothing | no |
| Bullying | Overt bullying victim items including: personal belongings stolen, threatened/blackmailed, hit/beaten up | weekly | no |
|  | Relational bullying victim items including: do something didn't want to, told lies about child | weekly | no |
|  | Friends tried to get teenager to do things didn't want to / told lies about teenager | weekly | no |
|  | Child has been bullied | all the time | no |
|  | Upset by name calling/exclusion from groups or bullying | Most days | no |
|  | Someone threatened/blackmailed teenager | weekly | no |
| Violence between parents | Physically cruel | yes, affected them | yes, the reported age can be retrospective |
|  | Kicked, bitten or hit each other | yes | no |
|  | Physically twisted arm | yes | no |
|  | Throw(n) bodily | yes | no |
|  | Beaten each other up | yes | no |
|  | Choke or strangle each other | yes | no |
|  | Threatened each other with knife | yes | no |
|  | Used knife or other weapon on each other | yes | no |
| Substance abuse in household | Smoked cannabis | every day | no |
|  | Hard drug use (including crack, heroin, amphetamine, opiate, cocaine, methadone, meth) | yes | no |
|  | Hard drug addiction | yes, recently | no |
|  | Alcoholism/ Drink problem | yes, ever / yes, saw doctor | no |
|  | Alcohol Use Disorders Identification Test (AUDIT) score | >8 | no |
| Parental mental health problems or attempted suicide | Parent has hurt themselves on purpose | yes | no |
|  | Parent has attempted suicide | yes | yes, the reported age can be retrospective |
|  | Taken medication for anxiety or depression | yes | no |
|  | Edinburgh Postnatal Depression Scale (EPDS) | >12 | no |
|  | Schizophrenia | yes, either current or ever | Both |
|  | Bulimia, anorexia nervosa | yes, recently | Both |
|  | Ever admitted to hospital for psychiatric or mental health problems | yes | yes |
| Parent convicted offence | Court conviction | yes | no |
|  | Convicted of an offence | yes | yes, the reported age can be retrospective |
| Parental separation | Parent reports divorce/separation | yes | no |
|  | Your parents have divorced/separated | yes | no |
|  | Parent still has the same partner/husband | no | no |

*Supplemental Table 2 Distributions of outcome and ACE variables in the imputation datasets and in observed data (i.e. without imputation) in boys and girls.*

|  | **Analysis 1: Education** | | | | | | | **Analysis 2: Health** | | | | | | |
| --- | --- | --- | --- | --- | --- | --- | --- | --- | --- | --- | --- | --- | --- | --- |
| **Variable** | **Boys** | | | **Girls** | | | **p-value** Gender difference imputed education data | **Boys** | | | **Girls** | | | **p-value** Gender difference imputed health data |
|  | **% data imputed** | **Distribution**  Mean (SE) for continuous variables  % for categorical variables  In | | **% data imputed** | **Distribution**  Mean (SE) for continuous variables  % for categorical variables  In | |  | **% data imputed** | **Distribution**  Mean (SE) for continuous variables  % for categorical variables  In | | **% data imputed** | **Distribution**  Mean (SE) for continuous variables  % for categorical variables  In | |  |
|  |  | imputed | observed |  | imputed | observed |  |  | imputed | observed |  | imputed | observed |  |
| **OUTCOME** | | | | | | | | | | | | | | |
| <5 GCSEs including math and English at grades A*-C | 0.2 | 50.7 | 50.6 | 0.1 | 40.3 | 40.3 | <0.01 | n/a | n/a | n/a | n/a | n/a | n/a | n/a |
| BMI-Z at age 17 | n/a | n/a | n/a | n/a | n/a | n/a | n/a | 1.8 | 0.38 (0.03) | 0.38 (0.03) | 1.8 | 0.4 (0.02) | 0.4 (0.02) | 0.68 |
| AUDIT score at age 17 | n/a | n/a | n/a | n/a | n/a | n/a | n/a | 19.8 | 8.64 (0.13) | 8.43 (0.14) | 19.9 | 8.12 (0.11) | 7.93 (0.12) | <0.01 |
| Obesity at age 17 | n/a | n/a | n/a | n/a | n/a | n/a | n/a | 1.8 | 6.3 | 6.3 | 1.8 | 8.0 | 8.0 | 0.02 |
| Depression at age 17 | n/a | n/a | n/a | n/a | n/a | n/a | n/a | 10.6 | 5.5 | 4.5 | 10.6 | 11.1 | 10.6 | <0.01 |
| Smoking at age 17 | n/a | n/a | n/a | n/a | n/a | n/a | n/a | 17.2 | 18.6 | 16.0 | 17.6 | 20.2 | 17.7 | 0.21 |
| Illicit drug use at age 17 | n/a | n/a | n/a | n/a | n/a | n/a | n/a | 18.8 | 17.9 | 15.2 | 19.3 | 14.6 | 12.5 | <0.01 |
| Harmful alcohol use at age 17 | n/a | n/a | n/a | n/a | n/a | n/a | n/a | 19.8 | 11.7 | 10.7 | 19.9 | 10.3 | 9.5 | 0.19 |
| **ADVERSE CHILDHOOD EXPERIENCES** | | | | | | | | | | | | | | |
| Categorical ACE-score 0 | 71.6 | 15.5 | 20.7 | 67.0 | 16.7 | 23.4 | 0.32 | 43.8 | 17.3 | 20.4 | 44.6 | 18.8 | 23.9 | 0.21 |
| 1 |  | 24.4 | 30 |  | 22.8 | 26.4 |  |  | 26.9 | 30.8 |  | 24.4 | 27.9 |  |
| 2 3 |  | 36.3 | 34.7 |  | 36.7 | 36.3 |  |  | 35.3 | 34.5 |  | 36.7 | 35.8 |  |
| 4+ |  | 23.8 | 14.6 |  | 23.8 | 13.9 |  |  | 20.5 | 14.3 |  | 20.1 | 12.4 |  |
| physical abuse | 48.8 | 17.0 | 13.2 | 44.0 | 20.9 | 16.6 | <0.01 | 28.6 | 20.9 | 17.2 | 26.8 | 20.7 | 17.5 | 0.9 |
| sexual abuse | 24.9 | 2.3 | 0.9 | 23.0 | 6 | 4.7 | <0.01 | 11.7 | 2.4 | 1.3 | 12.3 | 7.0 | 5.9 | <0.01 |
| emotional abuse | 44.0 | 24.0 | 18.8 | 40.8 | 23.7 | 19.5 | 0.8 | 25.7 | 22.2 | 17.8 | 25.7 | 23.3 | 19.9 | 0.5 |
| emotional neglect | 54.7 | 26.5 | 21.4 | 47.1 | 21.2 | 17.6 | <0.01 | 16.6 | 21.4 | 20.0 | 17.6 | 18.6 | 16.8 | 0.04 |
| bullying | 42.0 | 28.6 | 27.2 | 36.9 | 23.7 | 21.3 | <0.01 | 9.7 | 31.0 | 30.4 | 11.8 | 23.6 | 22.6 | <0.01 |
| violence between parents | 47.4 | 25.0 | 18.8 | 46.3 | 25.7 | 19.7 | 0.52 | 26.5 | 20.8 | 16.4 | 30.1 | 22.3 | 17.7 | 0.35 |
| substance household | 38.9 | 15.2 | 9.9 | 38.6 | 15.0 | 9.1 | 0.85 | 21.8 | 12.1 | 8.3 | 25.6 | 12.3 | 8.5 | 0.87 |
| mental health problems or suicide | 39.3 | 47.9 | 42.7 | 37.7 | 49.3 | 44.2 | 0.29 | 20.3 | 43.7 | 40.0 | 23.0 | 45.7 | 42.1 | 0.23 |
| parent convicted offence | 36.6 | 10.2 | 6.8 | 36.5 | 10.9 | 7.6 | 0.47 | 19.7 | 9.3 | 6.6 | 23.0 | 9.1 | 6.9 | 0.91 |
| parental separation | 46.1 | 33.0 | 24.3 | 44.4 | 34.7 | 27.2 | 0.23 | 26.3 | 27.3 | 21.3 | 29.0 | 29.0 | 24.1 | 0.3 |

*Supplemental Table 3 Description of the variables in the imputation model. Most variables were part of both the educational attainment as well as the health outcome imputation model, but outcome variables and certain auxiliary variables were specific to one of the two analyses (final column).*

| **Variable** | **Type of variable** | **Regression model to predict missing in this variable** | **How variable was entered when used to predict missing in other variables** | **Analysis variable was in the imputation model** |
| --- | --- | --- | --- | --- |
| **OUTCOME** | | | | |
| <5 GCSEs including math and English at grades A*-C | dichotomous | Logistic regression | dichotomous | Education |
| BMI-Z at age 17 | continuous | Predictive mean matching | continuous | Health |
| Obesity at age 17 | dichotomous | Passive imputation to split the BMI-Z score based on IOTF cut-offs for boys (>=2.288) and girls (>=2.192) | dichotomous^3^ | Health |
| Depression at age 17 | dichotomous | Logistic regression | dichotomous | Health |
| Smoking at age 17 | dichotomous | Logistic regression | dichotomous | Health |
| Illicit drug use at age 17 | dichotomous | Logistic regression | dichotomous | Health |
| AUDIT score at age 17 | continuous | Predictive mean matching | continuous | Health |
| Harmful alcohol use at age 17 | dichotomous | Passive imputation to split the AUDIT score (>=16) | dichotomous^3^ | Health |
| **ADVERSE CHILDHOOD EXPERIENCES** | | | | |
| Categorical ACE-score | categorical (4) | After passive imputation split the sum of 10 ACEs into 0,1,2-3 and 4+ ACEs^1^ | n/a^2^ | Both |
| physical abuse | dichotomous | Logistic regression | dichotomous | Both |
| sexual abuse | dichotomous | Logistic regression | dichotomous | Both |
| emotional abuse | dichotomous | Logistic regression | dichotomous | Both |
| emotional neglect | dichotomous | Logistic regression | dichotomous | Both |
| bullying | dichotomous | Logistic regression | dichotomous | Both |
| violence between parents | dichotomous | Logistic regression | dichotomous | Both |
| substance household | dichotomous | Logistic regression | dichotomous | Both |
| mental health problems or suicide | dichotomous | Logistic regression | dichotomous | Both |
| parent convicted offence | dichotomous | Logistic regression | dichotomous | Both |
| parental separation | dichotomous | Logistic regression | dichotomous | Both |
| **COVARIATES** | | | | |
| Gender | categorical (2) | n/a | categorical (2) | Both |
| Household social class at 18wks gestation | categorical (6) | Polytomous (unordered) regression | 5 indicator variables | Both |
| Ethnicity child | categorical (2) | Logistic regression | 1 indicator variables | Both |
| Maternal age in years at delivery | continuous | Predictive mean matching | continuous | Both |
| Home ownership mother during pregnancy | categorical (7) | Polytomous (unordered) regression | 6 indicator variables | Both |
| Marital status mother during pregnancy | categorical (6) | Polytomous (unordered) regression | 5 indicator variables | Both |
| Parity | continuous | Predictive mean matching | continuous | Both |
| Self-reported highest educational level mother | categorical (5) | Polytomous (unordered) regression | 4 indicator variables | Both |
| Mother-reported highest educational level partner | categorical (5) | Polytomous (unordered) regression | 4 indicator variables | Both |
| Maternal depression score (EPDS) at 18 wks gestation | continuous | Predictive mean matching | continuous | Both |
| Maternal depression score (EPDS) at 32 wks gestation | continuous | Predictive mean matching | continuous | Both |
| Partner depression score (EPDS) at 18 wks gestation | continuous | Predictive mean matching | continuous | Both |
| **AUXILIARY IMPUTATION** | | | | |
| social class | dichotomous | Logistic regression | dichotomous | Both |
| financial difficulties | dichotomous | Logistic regression | dichotomous | Both |
| satisfaction with neighbourhood | dichotomous | Logistic regression | dichotomous | Both |
| social support of child | dichotomous | Logistic regression | dichotomous | Both |
| social support of parent | dichotomous | Logistic regression | dichotomous | Both |
| violence between child and partner | dichotomous | Logistic regression | dichotomous | Both |
| physical illness of the child | dichotomous | Logistic regression | dichotomous | Both |
| physical illness of a parent | dichotomous | Logistic regression | dichotomous | Both |
| parent-child bond | dichotomous | Logistic regression | dichotomous | Both |
| Birthweight child in grams | continuous | Predictive mean matching | continuous | Both |
| Gestational age in weeks at delivery | continuous | Predictive mean matching | continuous | Both |
| Maternal pre-pregnancy weight (Kg) | continuous | Predictive mean matching | continuous | Both |
| Maternal pre-pregnancy BMI | continuous | Predictive mean matching | continuous | Both |
| Self-reported highest educational level partner | categorical (5) | Polytomous (unordered) regression | 4 indicator variables | Both |
| Partner-reported highest educational level mother | categorical (5) | Polytomous (unordered) regression | 4 indicator variables | Both |
| Mother s partner was emotionally cruel when child was 18yrs | dichotomous | Logistic regression | dichotomous | Both |
| Antidepressant use by mother when child was 18yrs | dichotomous | Logistic regression | dichotomous | Both |
| Maternal depression score (EPDS) when child was 18yrs | continuous | Predictive mean matching | continuous | Both |
| Mother separated from partner when child was 18yrs | dichotomous | Logistic regression | dichotomous | Both |
| Maternal AUDIT score when child was 18yrs | continuous | Predictive mean matching | continuous | Both |
| Paternal AUDIT score when child was 18yrs | continuous | Predictive mean matching | continuous | Both |
| Partner of child used physical force when child was 18-21yrs | dichotomous | Logistic regression | dichotomous | Both |
| Partner of child used more severe physical force when child was 18-21yrs | dichotomous | Logistic regression | dichotomous | Both |
| Partner of child have pressured them into kissing/touching when child was 18-21yrs | dichotomous | Logistic regression | dichotomous | Both |
| Partner of child physically forced them into kissing/touching when child was 18-21yrs | dichotomous | Logistic regression | dichotomous | Both |
| Partner of child used pressured them into sexual intercourse when child was 18-21yrs | dichotomous | Logistic regression | dichotomous | Both |
| Partner of child physically forced them into sexual intercourse when child was 18-21yrs | dichotomous | Logistic regression | dichotomous | Both |
| Partner of child made them feel scared of frightened when child was 18-21yrs | dichotomous | Logistic regression | dichotomous | Both |
| Maternal smoking during the 1st trimester of pregnancy | dichotomous | Logistic regression | dichotomous | Both |
| Maternal smoking during the 2nd trimester of pregnancy | dichotomous | Logistic regression | dichotomous | Both |
| Maternal smoking during the 3rd trimester of pregnancy (prospectively reported) | dichotomous | Logistic regression | dichotomous | Both |
| Maternal smoking during the 3rd trimester of pregnancy (retrospectively reported) | dichotomous | Logistic regression | dichotomous | Both |
| Mother became homeless during pregnancy | categorical (2) | Logistic regression | 1 indicator variables | Education |
| Difficulty affording food during pregnancy | categorical (4) | Polytomous (unordered) regression | 3 indicator variables | Education |
| Difficulty affording heating during pregnancy | categorical (4) | Polytomous (unordered) regression | 3 indicator variables | Education |
| Mother’s opinion of neighbourhood during pregnancy | categorical (4) | Polytomous (unordered) regression | 3 indicator variables | Education |
| Mother divorced since pregnancy | dichotomous | Logistic regression | dichotomous | Education |
| Partner hard drug use during pregnancy | categorical (2) | Logistic regression | 1 indicator variables | Education |
| Key stage 1: School year taken | categorical (3) | Polytomous (unordered) regression | 2 indicator variables | Education |
| Key stage 1: Summary score (prorated) | continuous | Predictive mean matching | continuous | Education |
| Key Stage 2: Total marks achieved in English test (sum of reading and writing tests) | continuous | Predictive mean matching | continuous | Education |
| Key Stage 2: Total marks achieved in Maths test (sum of Paper A, Paper B and mental arithmetic tests) | continuous | Predictive mean matching | continuous | Education |
| Key Stage 2: Total marks achieved in Science test (sum of Paper A and Paper B tests) | continuous | Predictive mean matching | continuous | Education |
| Key Stage 2: Total point score as used in the valued added calculations | continuous | Predictive mean matching | continuous | Education |
| Key Stage 3: Total marks achieved in English test (sum of reading and writing tests) | continuous | Predictive mean matching | continuous | Education |
| Key Stage 3: Total marks achieved in Maths test (sum of Paper A, Paper B and mental arithmetic tests) | continuous | Predictive mean matching | continuous | Education |
| Key Stage 3: Total marks achieved in Science test (sum of Paper A and Paper B tests) | continuous | Predictive mean matching | continuous | Education |
| Key Stage 3: Total point score as used in the valued added calculations | continuous | Predictive mean matching | continuous | Education |
| Key Stage 4: Deprivation Indicator - IDACI score (as used in CVA Model) | continuous | Predictive mean matching | continuous | Education |
| Key Stage 4: Is pupil known to be eligible for FSM? | dichotomous | Logistic regression | dichotomous | Education |
| Key Stage 4: Does pupil have SEN - Action Plus? | dichotomous | Logistic regression | dichotomous | Education |
| Key Stage 4: Does pupil have SEN - school action? | dichotomous | Logistic regression | dichotomous | Education |
| Key Stage 4: Total GCSE and equivalents new style point score | continuous | Predictive mean matching | continuous | Education |
| Key Stage 4: Total GCSE/GNVQ new style point score | continuous | Predictive mean matching | continuous | Education |
| >=5 GCSEs including math and English at grades A*-G | dichotomous | Logistic regression | dichotomous | Education |
| Key Stage 4: Achieved at least 1 GCSE or equivalent at grade A*-G | dichotomous | Logistic regression | dichotomous | Education |
| Key Stage 4: Achieved 5 or more GCSE/GNVQs at grades A*-C | dichotomous | Logistic regression | dichotomous | Education |
| Key Stage 4: Achieved 5 or more GCSE/GNVQs at grades A*-G | dichotomous | Logistic regression | dichotomous | Education |
| Key Stage 4: Number of Full GCSE qualifications at grades A*-C (GCSE equivalencies) | continuous | Predictive mean matching | continuous | Education |
| Key Stage 4: Number of Full GCSE qualifications at grades A*-G (GCSE equivalencies) | continuous | Predictive mean matching | continuous | Education |
| Key Stage 4: Capped GCSE and equivalents new style point | continuous | Predictive mean matching | continuous | Education |
| Key Stage 5: Participating at A levels | dichotomous | n/a | dichotomous | Education |
| Key Stage 5: Student achieved equivalent of 2 A levels | categorical (2) | Logistic regression | 1 indicator variables | Education |
| Key Stage 5: Total re-scaled point score of candidate s entries | continuous | Predictive mean matching | continuous | Education |
| Key Stage 5: Total GCE A Level and equivalent points score based on new QCA points | continuous | Predictive mean matching | continuous | Education |
| Key Stage 5: Total number of GCE/VCE A/AS Level & GCE AS/VCE Double Award Level passes (A Levels) | continuous | Predictive mean matching | continuous | Education |
| BMI at age 9 | continuous | Predictive mean matching | continuous | Health |
| Maternal smoking at age 17y | dichotomous | Logistic regression | dichotomous | Health |
| Smoker at age 13 | dichotomous | Logistic regression | dichotomous | Health |
| Smoker at age 15.5 | dichotomous | Logistic regression | dichotomous | Health |
| Smoker at age 16 | dichotomous | Logistic regression | dichotomous | Health |
| Smoker at age 18 | dichotomous | Logistic regression | dichotomous | Health |
| AUDIT score at age 16 | continuous | Predictive mean matching | continuous | Health |
| AUDIT score at age 18 | continuous | Predictive mean matching | continuous | Health |
| Alcohol use at age 13 | dichotomous | Logistic regression | dichotomous | Health |
| MFQ score at age 10.5 | continuous | Predictive mean matching | continuous | Health |
| MFQ score at age 12.5 | continuous | Predictive mean matching | continuous | Health |
| MFQ score at age 16 | continuous | Predictive mean matching | continuous | Health |
| MFQ score at age 17 | continuous | Predictive mean matching | continuous | Health |
| MFQ score at age 18 | continuous | Predictive mean matching | continuous | Health |
| Maternal MFQ score at age 16 | continuous | Predictive mean matching | continuous | Health |
| Illicit drug use at age 13 | dichotomous | Logistic regression | dichotomous | Health |
| Illicit drug use at age 14 | dichotomous | Logistic regression | dichotomous | Health |
| Illicit drug use at age 15.5 | dichotomous | Logistic regression | dichotomous | Health |
| Illicit drug use at age 20 | dichotomous | Logistic regression | dichotomous | Health |
| Illicit drug use at age 16 | dichotomous | Logistic regression | dichotomous | Health |
| Illicit drug use at age 18 | dichotomous | Logistic regression | dichotomous | Health |
| Cannabis use at age 15.5 | dichotomous | Logistic regression | dichotomous | Health |
| Cannabis use at age 16 | dichotomous | Logistic regression | dichotomous | Health |
| Cannabis use at age 18 | dichotomous | Logistic regression | dichotomous | Health |
| Cannabis use at age 20 | dichotomous | Logistic regression | dichotomous | Health |

^1^Formula used for the passive imputation of the ACE score: ~I(as.integer(emotional_abuse)+as.integer(physical_abuse)+as.integer(sexual_abuse)+as.integer(mental_suicide_household)+as.integer(parental_separation)+as.integer(emotional_neglect)+as.integer(violence_household)+as.integer(bullying)+as.integer(substance_household)+as.integer(parent_convicted))

^2^ None of the passively imputed variables (ACE count score variable, obesity and harmful alcohol use) were used as a predictor of missingness for the other variables in the imputation model.

*Supplemental table 4 Distributions of imputed characteristics in the imputation datasets and in observed data (i.e. without imputation).*

| **Variable** | **Analysis 1: Education** | | | **Analysis 2: Health** | | |
| --- | --- | --- | --- | --- | --- | --- |
|  | **% data imputed** | **Distribution**  Mean (SE) for continuous variables  % for categorical variables  In | | **% data imputed** | **Distribution** Mean (SE) for continuous variables  % for categorical variables  In | |
|  |  | imputed | observed |  | imputed | observed |
| **OUTCOME** | | | | | | |
| <5 GCSEs including math and English at grades A*-C | 0.2 | 46.5 | 46.5 | n/a | n/a | n/a |
| BMI-Z at age 17 | n/a | n/a | n/a | 1.8 | 0.39 (0.02) | 0.39 (0.02) |
| AUDIT score at age 17 | n/a | n/a | n/a | 19.9 | 8.35 (0.08) | 8.15 (0.09) |
| Obesity at age 17 | n/a | n/a | n/a | 1.8 | 7.3 | 7.2 |
| Depression at age 17 | n/a | n/a | n/a | 10.6 | 8.7 | 7.9 |
| Smoking at age 17 | n/a | n/a | n/a | 17.4 | 19.5 | 16.9 |
| Illicit drug use at age 17 | n/a | n/a | n/a | 19.1 | 16.1 | 13.7 |
| Harmful alcohol use at age 17 | n/a | n/a | n/a | 19.9 | 10.9 | 10 |
| **ADVERSE CHILDHOOD EXPERIENCES** | | | | | | |
| Categorical ACE-score | 69.3 |  |  | 44.3 |  |  |
| 0 |  | 16.1 | 22.1 |  | 18.2 | 22.4 |
| 1 |  | 23.6 | 28.1 |  | 25.5 | 29.1 |
| 2 3 |  | 36.5 | 35.5 |  | 36.1 | 35.2 |
| 4+ |  | 23.8 | 14.3 |  | 20.3 | 13.2 |
| physical abuse | 46.4 | 19 | 15 | 27.6 | 20.8 | 17.4 |
| sexual abuse | 23.9 | 4.1 | 2.8 | 12 | 5 | 3.9 |
| emotional abuse | 42.4 | 23.9 | 19.2 | 25.7 | 22.8 | 19 |
| emotional neglect | 50.9 | 23.9 | 19.3 | 17.2 | 19.8 | 18.2 |
| bullying | 39.5 | 26.2 | 24.2 | 10.9 | 26.9 | 26 |
| violence between parents | 46.8 | 25.3 | 19.3 | 28.5 | 21.6 | 17.1 |
| substance household | 38.7 | 15.1 | 9.5 | 24 | 12.2 | 8.4 |
| mental health problems or suicide | 38.5 | 48.6 | 43.4 | 21.8 | 44.8 | 41.1 |
| parent convicted offence | 36.5 | 10.5 | 7.2 | 21.6 | 9.2 | 6.7 |
| parental separation | 45.2 | 33.8 | 25.8 | 27.8 | 28.3 | 22.8 |
| **COVARIATES** *basic model* | | | | | | |
| Gender (Male) | 0 | 49.6 | 49.6 | 0 | 56 | 56 |
| **COVARIATES** *adjusted model* | | | | | | |
| Ethnicity child (non-white) | 9.9 | 5 | 4.1 | 8.3 | 5.3 | 4.3 |
| Household social class at 18wks gestation | 10.5 |  |  | 9 |  |  |
| I - Professional |  | 8.4 | 8.2 |  | 12.6 | 12.9 |
| II - Managerial and technical |  | 36.4 | 37.5 |  | 41.2 | 42.7 |
| IIINM - Skilled non-manual |  | 31.3 | 32.5 |  | 28.3 | 28.5 |
| IIIM - Skilled manual |  | 13.7 | 13.3 |  | 10.7 | 10 |
| IV - Partly skilled |  | 8 | 6.9 |  | 5.9 | 5.1 |
| V - Unskilled |  | 2.3 | 1.6 |  | 1.2 | 0.8 |
| Home ownership mother during pregnancy | 6.8 |  |  | 6.7 |  |  |
| Mortgaged |  | 73.3 | 75.3 |  | 80.7 | 82.5 |
| Owned |  | 2.3 | 2.1 |  | 2.3 | 2.2 |
| Council rented |  | 12.7 | 12.2 |  | 8 | 7.3 |
| Furnished private rental |  | 3.8 | 3.4 |  | 2.9 | 2.6 |
| Unfurnished private rental |  | 2.9 | 2.6 |  | 2 | 1.8 |
| Housing authority rented |  | 1.8 | 1.4 |  | 1 | 0.8 |
| Other |  | 3.2 | 3 |  | 3 | 2.8 |
| Marital status mother during pregnancy | 6.3 |  |  | 6 |  |  |
| Never married |  | 18.4 | 17.2 |  | 14.6 | 13.7 |
| Widowed |  | 0.2 | 0.1 |  | 0.1 | 0.1 |
| Divorced |  | 4.1 | 3.9 |  | 3.7 | 3.5 |
| Separated |  | 1.7 | 1.4 |  | 1.4 | 1.2 |
| 1st marriage |  | 69.2 | 71.1 |  | 73.9 | 75.4 |
| Marriage 2 or 3 |  | 6.4 | 6.4 |  | 6.2 | 6.1 |
| Self-reported highest educational level mother | 7.9 |  |  | 6.9 |  |  |
| Certificate of Secondary Education (CSE) |  | 19.7 | 18.9 |  | 12.2 | 11.4 |
| Vocational |  | 10.6 | 10.2 |  | 7.8 | 7.4 |
| O level |  | 35.4 | 36.7 |  | 33.8 | 34.3 |
| A level |  | 22.2 | 22.7 |  | 27.4 | 28 |
| Degree |  | 12.1 | 11.5 |  | 18.8 | 18.8 |
| Mother-reported highest educational level partner | 11.3 |  |  | 9.2 |  |  |
| CSE |  | 27 | 25.2 |  | 19.9 | 17.9 |
| Vocational |  | 9.2 | 8.9 |  | 7.7 | 7.4 |
| O level |  | 21.9 | 22.5 |  | 20.9 | 21.3 |
| A level |  | 25.3 | 26.8 |  | 27.6 | 28.6 |
| Degree |  | 16.5 | 16.5 |  | 24 | 24.8 |
| Maternal age in years at delivery | 3.3 | 28.23 (0.05) | 28.26 (0.05) | 4.3 | 29.23 (0.07) | 29.27 (0.07) |
| Parity | 7.2 | 0.83 (0.01) | 0.83 (0.01) | 7.1 | 0.75 (0.01) | 0.74 (0.01) |
| Maternal depression score (EPDS) at 18 wks gestation | 13.3 | 6.99 (0.05) | 6.79 (0.05) | 12.5 | 6.49 (0.07) | 6.36 (0.07) |
| Maternal depression score (EPDS) at 32 wks gestation | 10.8 | 7.11 (0.05) | 6.91 (0.05) | 9.6 | 6.66 (0.07) | 6.53 (0.07) |
| Partner depression score (EPDS) at 18 wks gestation | 28 | 4.36 (0.04) | 4.12 (0.05) | 24 | 4.25 (0.06) | 4.03 (0.06) |
| **IMPUTATION VARIABLES** | | | | | | |
| social class | 52.5 | 13.3 | 10.1 | 52.2 | 13.5 | 7.7 |
| financial difficulties | 36.7 | 19.6 | 13.8 | 22.3 | 15.3 | 11.1 |
| satisfaction with neighbourhood | 26.3 | 11.5 | 9.2 | 7.8 | 11.4 | 10.4 |
| social support of child | 42.5 | 13.7 | 10.5 | 10.8 | 11.3 | 10.4 |
| social support of parent | 34 | 14.6 | 11.1 | 20.3 | 13 | 10.4 |
| violence between child and partner | 65 | 16.7 | 11 | 46.7 | 13.4 | 10 |
| physical illness of the child | 21.9 | 10 | 8.7 | 12.3 | 8.9 | 8 |
| physical illness of a parent | 51.2 | 28 | 24 | 34.1 | 27 | 24.3 |
| parent-child bond | 43.2 | 23.7 | 19.3 | 24.9 | 22.7 | 19.5 |
| Self-reported highest educational level partner | 27.7 |  |  | 23.5 |  |  |
| CSE |  | 23.9 | 20.8 |  | 17.9 | 14.7 |
| Vocational |  | 10.2 | 9 |  | 8.1 | 6.7 |
| O level |  | 22.4 | 23.7 |  | 22 | 22.2 |
| A level |  | 26.2 | 28.6 |  | 27.6 | 29.6 |
| Degree |  | 17.3 | 17.8 |  | 24.4 | 26.8 |
| Partner-reported highest educational level mother | 29.4 |  |  | 25 |  |  |
| CSE |  | 22.2 | 19.6 |  | 14.9 | 11.8 |
| Vocational |  | 10.5 | 9.8 |  | 8.3 | 7.2 |
| O level |  | 32.7 | 34.7 |  | 30.8 | 31.4 |
| A level |  | 21.7 | 22.9 |  | 26.1 | 28.4 |
| Degree |  | 12.9 | 13 |  | 19.9 | 21.1 |
| Birthweight child in grams | 4.4 | 3412.03 (5.5) | 3416.83 (5.58) | 5.3 | 3416.03 (7.67) | 3419.3 (7.84) |
| Gestational age in weeks at delivery | 3.3 | 39.48 (0.02) | 39.49 (0.02) | 4.3 | 39.47 (0.03) | 39.48 (0.03) |
| Maternal pre-pregnancy weight (Kg) | 13.2 | 61.94 (0.11) | 61.92 (0.12) | 11.7 | 61.69 (0.15) | 61.63 (0.16) |
| Maternal pre-pregnancy BMI | 14 | 23.05 (0.04) | 23.04 (0.04) | 12.5 | 22.86 (0.05) | 22.83 (0.06) |
| Maternal smoking during the 1st trimester of pregnancy | 5.8 | 24.7 | 22.8 | 5.9 | 17.1 | 16.1 |
| Maternal smoking during the 2nd trimester of pregnancy | 5.8 | 20 | 18.3 | 5.9 | 13.5 | 12.7 |
| Maternal smoking during the 3rd trimester of pregnancy (prospectively reported) | 15.8 | 21.4 | 19.4 | 14.7 | 13.8 | 13.5 |
| Maternal smoking during the 3rd trimester of pregnancy (retrospectively reported) | 9.5 | 21.1 | 18.5 | 8.6 | 13.8 | 12.6 |
| Mother became homeless during pregnancy | 11.9 | 3 | 2.1 | n/a | n/a | n/a |
| Mother divorced since pregnancy | 13.8 | 4.7 | 3.3 | n/a | n/a | n/a |
| Partner hard drug use during pregnancy | 29.1 | 4.1 | 1.7 | n/a | n/a | n/a |
| Mother's partner was emotionally cruel when child was 18yrs | 65.9 | 16.4 | 4.4 | 41.4 | 11 | 4.1 |
| Antidepressant use by mother when child was 18yrs | 67.5 | 23.7 | 11.2 | 44.1 | 16.6 | 10.3 |
| Mother separated from partner when child was 18yrs | 65.9 | 18 | 4.6 | 41.4 | 11.2 | 3.9 |
| Partner of child used physical force when child was 18-21yrs | 72.9 | 52.6 | 19.3 | 51.2 | 46.4 | 18 |
| Partner of child used more severe physical force when child was 18-21yrs | 72.9 | 47.4 | 13 | 51.3 | 41.8 | 11.8 |
| Partner of child have pressured them into kissing/touching when child was 18-21yrs | 72.9 | 47 | 14.1 | 51.2 | 42.2 | 13.2 |
| Partner of child physically forced them into kissing/touching when child was 18-21yrs | 73 | 46 | 11.3 | 51.3 | 40.4 | 10.2 |
| Partner of child used pressured them into sexual intercourse when child was 18-21yrs | 73 | 47.6 | 15.1 | 51.4 | 43.7 | 14.3 |
| Partner of child physically forced them into sexual intercourse when child was 18-21yrs | 73.1 | 45.7 | 10.8 | 51.5 | 41.1 | 10 |
| Partner of child made them feel scared of frightened when child was 18-21yrs | 73.2 | 49.3 | 19.1 | 51.7 | 45.6 | 17.9 |
| Difficulty affording food during pregnancy | 10.4 |  |  | n/a | n/a | n/a |
| Not difficult |  | 73.7 | 76.5 | n/a | n/a | n/a |
| Some difficulty |  | 16.4 | 15.6 | n/a | n/a | n/a |
| Fairly difficult |  | 8 | 6.7 | n/a | n/a | n/a |
| Very difficult |  | 1.9 | 1.2 | n/a | n/a | n/a |
| Difficulty affording heating during pregnancy | 10.4 |  |  | n/a | n/a | n/a |
| Not difficult |  | 69.5 | 72.4 | n/a | n/a | n/a |
| Some difficulty |  | 17.7 | 17.3 | n/a | n/a | n/a |
| Fairly difficult |  | 9.1 | 7.7 | n/a | n/a | n/a |
| Very difficult |  | 3.7 | 2.6 | n/a | n/a | n/a |
| Mother's opinion of neighbourhood during pregnancy | 8.2 | 41 | 41.7 | n/a | n/a | n/a |
| Very good area |  | 41 | 41.7 | n/a | n/a | n/a |
| Fairly good area |  | 51.2 | 51.3 | n/a | n/a | n/a |
| Not very good area |  | 5.8 | 5.3 | n/a | n/a | n/a |
| Bad area |  | 2 | 1.6 | n/a | n/a | n/a |
| Maternal depression score (EPDS) when child was 18yrs | 66.2 | 8.92 (0.06) | 7.52 (0.09) | 42.1 | 7.91 (0.08) | 7.25 (0.1) |
| Maternal AUDIT score when child was 18yrs | 71.5 | 8.63 (0.04) | 8.01 (0.06) | 50 | 8.34 (0.05) | 8.04 (0.06) |
| Paternal AUDIT score when child was 18yrs | 84.7 | 9.75 (0.04) | 9.13 (0.08) | 69.1 | 9.38 (0.05) | 9.13 (0.08) |
| Key stage 1: School year taken | 12.4 |  |  | n/a | n/a | n/a |
| 1997 / 1998 |  | 21.1 | 20.9 | n/a | n/a | n/a |
| 1998 / 1999 |  | 60.7 | 60.8 | n/a | n/a | n/a |
| 1999 / 2000 |  | 18.2 | 18.3 | n/a | n/a | n/a |
| Key stage 1: Summary score (prorated) | 12.7 | 9.62 (0.04) | 9.53 (0.04) | n/a | n/a | n/a |
| Key Stage 2: Total marks achieved in English test (sum of reading and writing tests) | 4.4 | 58.15 (0.16) | 58.98 (0.16) | n/a | n/a | n/a |
| Key Stage 2: Total marks achieved in Maths test (sum of Paper A, Paper B and mental arithmetic tests) | 4.1 | 64.44 (0.22) | 65.42 (0.21) | n/a | n/a | n/a |
| Key Stage 2: Total marks achieved in Science test (sum of Paper A and Paper B tests) | 3.2 | 59.07 (0.12) | 59.48 (0.12) | n/a | n/a | n/a |
| Key Stage 2: Total point score as used in the valued added calculations | 82.1 | 84.86 (0.13) | 86.49 (0.31) | n/a | n/a | n/a |
| Key Stage 3: Total marks achieved in English test (sum of reading and writing tests) | 14.4 | 45.71 (0.18) | 46.84 (0.19) | n/a | n/a | n/a |
| Key Stage 3: Total marks achieved in Maths test (sum of Paper A, Paper B and mental arithmetic tests) | 12.3 | 81.91 (0.23) | 82.7 (0.23) | n/a | n/a | n/a |
| Key Stage 3: Total marks achieved in Science test (sum of Paper A and Paper B tests) | 11.8 | 97.24 (0.26) | 98.27 (0.26) | n/a | n/a | n/a |
| Key Stage 3: Total point score as used in the valued added calculations | 9.3 | 106.16 (0.24) | 105.57 (0.25) | n/a | n/a | n/a |
| Key Stage 4: Deprivation Indicator - IDACI score (as used in CVA Model) | 1.5 | 0.15 (0) | 0.15 (0) | n/a | n/a | n/a |
| Key Stage 4: Total GCSE and equivalents new style point score | 0.7 | 407.21 (1.5) | 409.18 (1.49) | n/a | n/a | n/a |
| Key Stage 4: Total GCSE/GNVQ new style point score | 0.2 | 357.83 (1.49) | 357.93 (1.49) | n/a | n/a | n/a |
| Key Stage 4: Number of Full GCSE qualifications at grades A*-C (GCSE equivalencies) | 0.2 | 5.61 (0.04) | 5.62 (0.04) | n/a | n/a | n/a |
| Key Stage 4: Number of Full GCSE qualifications at grades A*-G (GCSE equivalencies) | 0.2 | 7.86 (0.02) | 7.86 (0.02) | n/a | n/a | n/a |
| Key Stage 4: Is pupil known to be eligible for FSM? | 5.6 | 5.2 | 5.1 | n/a | n/a | n/a |
| Key Stage 4: Does pupil have SEN - Action Plus? | 7.4 | 5.6 | 4.8 | n/a | n/a | n/a |
| Key Stage 4: Does pupil have SEN - school action? | 7.4 | 8.2 | 8 | n/a | n/a | n/a |
| >=5 GCSEs including math and English at grades A*-G | 0.2 | 90.6 | 90.7 | n/a | n/a | n/a |
| Key Stage 4: Achieved at least 1 GCSE or equivalent at grade A*-G | 0.2 | 98.4 | 98.4 | n/a | n/a | n/a |
| Key Stage 4: Achieved 5 or more GCSE/GNVQs at grades A*-C | 0.2 | 63.7 | 63.7 | n/a | n/a | n/a |
| Key Stage 4: Achieved 5 or more GCSE/GNVQs at grades A*-G | 0.2 | 92 | 92.1 | n/a | n/a | n/a |
| Participating at A levels (Key Stage 5) | 0 | 58.5 | 58.5 | n/a | n/a | n/a |
| Capped GCSE and equivalents new style point | 0.9 | 322.13 (0.95) | 324.52 (0.92) | n/a | n/a | n/a |
| Key Stage 5: Student achieved equivalent of 2 A levels | 41.5 | 75.3 | 95.6 | n/a | n/a | n/a |
| Key Stage 5: Total re-scaled point score of candidate's entries | 55 | 250.48 (1.7) | 343.69 (2.42) | n/a | n/a | n/a |
| Key Stage 5: Total GCE A Level and equivalent points score based on new QCA points | 41.5 | 611.33 (2.95) | 752.97 (3.3) | n/a | n/a | n/a |
| Key Stage 5: Total number of GCE/VCE A/AS Level & GCE AS/VCE Double Award Level passes (A Levels) | 45.1 | 1.91 (0.02) | 2.8 (0.02) | n/a | n/a | n/a |
| BMI at age 9 | n/a | n/a | n/a | 47.7 | 17.52 (0.04) | 17.43 (0.06) |
| AUDIT score at age 16 | n/a | n/a | n/a | 34.9 | 6.98 (0.08) | 6.59 (0.09) |
| AUDIT score at age 18 | n/a | n/a | n/a | 52.3 | 10.12 (0.09) | 9.1 (0.12) |
| MFQ score at age 10.5 | n/a | n/a | n/a | 12 | 4.05 (0.05) | 3.97 (0.05) |
| MFQ score at age 12.5 | n/a | n/a | n/a | 13.4 | 4.13 (0.06) | 4.02 (0.06) |
| MFQ score at age 16 | n/a | n/a | n/a | 30.2 | 6.04 (0.08) | 5.8 (0.09) |
| MFQ score at age 17 | n/a | n/a | n/a | 14.8 | 6.67 (0.08) | 6.56 (0.08) |
| MFQ score at age 18 | n/a | n/a | n/a | 49.6 | 6.89 (0.09) | 6.62 (0.12) |
| MFQ score at age 16 mum | n/a | n/a | n/a | 28.2 | 2.28 (0.05) | 2.05 (0.05) |
| Maternal smoking at age 17y | n/a | n/a | n/a | 41.2 | 16.8 | 9.5 |
| Smoker at age 13 | n/a | n/a | n/a | 15.8 | 4.2 | 1.7 |
| Smoker at age 15.5 | n/a | n/a | n/a | 19.4 | 12 | 8.5 |
| Smoker at age 16 | n/a | n/a | n/a | 29.3 | 18.2 | 11.2 |
| Smoker at age 18 | n/a | n/a | n/a | 49.4 | 32.2 | 14.6 |
| Alcohol use at age 13 | n/a | n/a | n/a | 32.3 | 8.3 | 4.9 |
| Illicit drug use at age 13 | n/a | n/a | n/a | 13 | 3.4 | 1.8 |
| Illicit drug use at age 14 | n/a | n/a | n/a | 16 | 7.4 | 4.3 |
| Illicit drug use at age 15.5 | n/a | n/a | n/a | 25.1 | 16 | 12.4 |
| Illicit drug use at age 20 | n/a | n/a | n/a | 41.5 | 25.2 | 19.8 |
| Illicit drug use at age 16 | n/a | n/a | n/a | 33 | 22 | 15.2 |
| Illicit drug use at age 18 | n/a | n/a | n/a | 50.4 | 27.5 | 11.7 |
| Cannabis use at age 15.5 | n/a | n/a | n/a | 20.6 | 5.5 | 2.6 |
| Cannabis use at age 16 | n/a | n/a | n/a | 29.5 | 8.9 | 3 |
| Cannabis use at age 18 | n/a | n/a | n/a | 50.9 | 19.9 | 2.9 |
| Cannabis use at age 20 | n/a | n/a | n/a | 40.9 | 12.3 | 5.1 |

*Supplemental Table 5 Adjusted odds ratios (AOR) and 95% confidence intervals for the association between the ACE measures and educational attainment or health. The basic model was adjusted for gender, whereas the adjusted model also includes sociodemographic indicators. The p-value of the interaction test is for the comparison of the original model with a model including a gender interaction. Furthermore, model estimates of gender stratified analyses are given.*

| **Adversity** | **Model** | **Analysis 1: Education** | **Analysis 2: Health** | | | | |
| --- | --- | --- | --- | --- | --- | --- | --- |
|  |  | <5 good GCSE  AOR 95% CI | Depression  AOR 95% CI | Harmful alcohol use  AOR 95% CI | Illicit drug use AOR 95% CI | Obesity AOR 95% CI | Smoking AOR 95% CI |
| **Categorical ACE score** | Basic 1 ACE | 1.38(1.17,1.62)* | 1.38 (0.88,2.18) | 1.04 (0.74,1.46) | 1.45 (1.05,2)* | 1.39 (0.91,2.11) | 1.22 (0.91,1.65) |
|  | Basic 2-3 ACEs | 1.77(1.52,2.06)* | 2.17 (1.46,3.21)* | 1.32 (0.97,1.8) | 1.9 (1.42,2.54)* | 1.61 (1.1,2.35)* | 1.75 (1.35,2.27)* |
|  | Basic 4+ ACEs | 3.18(2.69,3.75)* | 3.15 (2.08,4.78)* | 1.59 (1.13,2.24)* | 3.31 (2.43,4.51)* | 1.81 (1.19,2.73)* | 2.78 (2.1,3.67)* |
|  | Adjusted 1 ACE | 1.37(1.15,1.63)* | 1.33 (0.84,2.09) | 1 (0.71,1.4) | 1.43 (1.04,1.98)* | 1.34 (0.88,2.05) | 1.19 (0.88,1.6) |
|  | Adjusted 2-3 ACEs | 1.57(1.33,1.86)* | 1.97 (1.31,2.96)* | 1.21 (0.88,1.67) | 1.8 (1.33,2.44)* | 1.48 (1,2.19) | 1.64 (1.25,2.14)* |
|  | Adjusted 4+ ACEs | 2(1.65,2.43)* | 2.49 (1.57,3.94)* | 1.37 (0.94,1.99) | 2.98 (2.13,4.18)* | 1.36 (0.86,2.16) | 2.31 (1.7,3.15)* |
|  | Adjusted complete cases 1 ACE | 1.21(0.92,1.61) | 1.01 (0.57,1.82) | 0.97 (0.61,1.55) | 1.71 (1.08,2.75)* | 0.93 (0.55,1.57) | 1.01 (0.67,1.53) |
|  | Adjusted complete cases 2-3 ACEs | 1.25(0.95,1.64) | 1.96 (1.19,3.34)* | 1.21 (0.78,1.9) | 2.3 (1.49,3.64)* | 1.32 (0.82,2.17) | 1.58 (1.09,2.32)* |
|  | Adjusted complete cases 4+ ACEs | 1.46(1.04,2.05)* | 2.24 (1.21,4.19)* | 0.98 (0.53,1.79) | 3.38 (2.01,5.77)* | 1.08 (0.56,2.06) | 2.11 (1.33,3.36)* |
|  | Gender interaction | 0.928 | 0.674 | 0.34 | 0.577 | 0.19 | 0.538 |
|  | Boys 1 ACE | 1.35(1.06,1.72)* | 1 (0.42,2.4) | 1.35 (0.79,2.31) | 1.37 (0.85,2.2) | 0.84 (0.45,1.56) | 1.13 (0.72,1.78) |
|  | Boys 2-3 ACEs | 1.59(1.26,2.01)* | 1.87 (0.86,4.08) | 1.66 (1,2.74)* | 1.83 (1.17,2.85)* | 1.02 (0.57,1.83) | 1.69 (1.12,2.55)* |
|  | Boys 4+ ACEs | 1.93(1.46,2.54)* | 1.78 (0.75,4.23) | 1.67 (0.91,3.05) | 2.72 (1.64,4.52)* | 1 (0.5,2) | 2.07 (1.27,3.37)* |
|  | Girls 1 ACE | 1.4(1.08,1.81)* | 1.49 (0.88,2.53) | 0.77 (0.48,1.23) | 1.5 (0.95,2.36) | 1.88 (1.04,3.37)* | 1.22 (0.81,1.83) |
|  | Girls 2-3 ACEs | 1.56(1.22,2)* | 2.01 (1.23,3.26)* | 0.96 (0.63,1.47) | 1.79 (1.14,2.79)* | 1.96 (1.13,3.39)* | 1.59 (1.1,2.28)* |
|  | Girls 4+ ACEs | 2.1(1.59,2.77)* | 2.89 (1.68,4.97)* | 1.18 (0.71,1.96) | 3.27 (2.03,5.29)* | 1.71 (0.9,3.23) | 2.42 (1.62,3.62)* |
| **Physical abuse** | Basic | 1.15(0.98,1.34) | 1.93 (1.46,2.55)* | 1.28 (0.99,1.67) | 1.63 (1.28,2.08)* | 1.42 (1.07,1.9)* | 1.79 (1.44,2.21)* |
|  | Adjusted | 0.91(0.76,1.07) | 1.83 (1.36,2.46)* | 1.22 (0.93,1.6) | 1.56 (1.21,2.02)* | 1.31 (0.97,1.77) | 1.68 (1.34,2.11)* |
|  | Adjusted complete cases | 0.83(0.67,1.01) | 2.07 (1.43,2.96)* | 1.01 (0.68,1.46) | 1.56 (1.14,2.11)* | 1.33 (0.89,1.96) | 1.68 (1.26,2.23)* |
|  | Gender interaction | 1 | 0.433 | 0.615 | 0.41 | 0.894 | 0.082 |
|  | Boys | 0.91(0.71,1.17) | 1.59 (0.89,2.83) | 1.13 (0.72,1.78) | 1.47 (1.01,2.15)* | 1.4 (0.86,2.27) | 1.41 (0.99,2.02) |
|  | Girls | 0.91(0.71,1.15) | 2.01 (1.4,2.89)* | 1.28 (0.89,1.84) | 1.68 (1.17,2.4)* | 1.24 (0.84,1.83) | 1.97 (1.47,2.65)* |
| **Sexual abuse** | Basic | 1.83(1.4,2.39)* | 2.26 (1.52,3.36)* | 1.54 (0.99,2.39) | 1.54 (1.01,2.34)* | 1.68 (1.05,2.69)* | 1.93 (1.37,2.73)* |
|  | Adjusted | 1.35(0.99,1.84) | 2.06 (1.35,3.13)* | 1.45 (0.92,2.3) | 1.42 (0.92,2.19) | 1.3 (0.8,2.12) | 1.61 (1.12,2.32)* |
|  | Adjusted complete cases | 1.25(0.83,1.87) | 2.5 (1.38,4.34)* | 2.04 (1.04,3.74)* | 1.79 (0.97,3.13)* | 1.19 (0.56,2.27) | 1.81 (1.07,2.98)* |
|  | Gender interaction | 0.188 | 0.393 | 0.846 | 0.129 | 0.467 | 0.722 |
|  | Boys | 1.95(0.98,3.89) | 3.39 (1.12,10.26)* | 1.3 (0.46,3.69) | 0.75 (0.24,2.35) | 2.3 (0.77,6.87) | 2.07 (0.95,4.52) |
|  | Girls | 1.2(0.85,1.68) | 1.94 (1.21,3.09)* | 1.49 (0.88,2.5) | 1.71 (1.06,2.75)* | 1.08 (0.61,1.91) | 1.47 (0.97,2.25) |
| **Emotional abuse** | Basic | 1.36(1.2,1.54)* | 1.64 (1.25,2.15)* | 1.21 (0.93,1.57) | 1.79 (1.44,2.24)* | 1.07 (0.78,1.46) | 1.42 (1.15,1.75)* |
|  | Adjusted | 1.14(0.98,1.31) | 1.41 (1.05,1.9)* | 1.11 (0.84,1.47) | 1.61 (1.27,2.05)* | 0.96 (0.68,1.34) | 1.28 (1.02,1.61)* |
|  | Adjusted complete cases | 1.12(0.94,1.33) | 1.3 (0.89,1.87) | 0.96 (0.66,1.38) | 1.82 (1.36,2.43)* | 1 (0.66,1.48) | 1.33 (1,1.76)* |
|  | Gender interaction | 0.966 | 0.388 | 0.977 | 0.955 | 0.756 | 0.969 |
|  | Boys | 1.14(0.93,1.4) | 1.08 (0.59,1.98) | 1.17 (0.77,1.79) | 1.71 (1.21,2.41)* | 1.13 (0.67,1.92) | 1.31 (0.93,1.85) |
|  | Girls | 1.15(0.93,1.42) | 1.55 (1.12,2.16)* | 1.1 (0.76,1.59) | 1.57 (1.13,2.17)* | 0.82 (0.53,1.27) | 1.22 (0.91,1.64) |
| **Emotional neglect** | Basic | 2.31(2.01,2.65)* | 1.23 (0.93,1.63) | 0.94 (0.72,1.24) | 1.09 (0.84,1.41) | 1.48 (1.11,1.97)* | 1.16 (0.93,1.45) |
|  | Adjusted | 1.9(1.64,2.2)* | 1.14 (0.85,1.52) | 0.92 (0.69,1.23) | 1.06 (0.81,1.39) | 1.24 (0.92,1.66) | 1.05 (0.83,1.32) |
|  | Adjusted complete cases | 1.81(1.49,2.2)* | 1.34 (0.9,1.95) | 1.08 (0.72,1.58) | 0.86 (0.6,1.21) | 1.37 (0.93,1.97) | 0.98 (0.71,1.32) |
|  | Gender interaction | 0.896 | 0.203 | 0.674 | 0.188 | 0.491 | 0.046 |
|  | Boys | 1.9(1.54,2.35)* | 1.44 (0.86,2.39) | 0.99 (0.67,1.47) | 0.89 (0.62,1.27) | 1.24 (0.79,1.95) | 0.79 (0.54,1.16) |
|  | Girls | 1.92(1.54,2.39)* | 1.02 (0.71,1.48) | 0.85 (0.55,1.29) | 1.24 (0.84,1.82) | 1.3 (0.88,1.91) | 1.27 (0.94,1.71) |
| **Bullying** | Basic | 1.52(1.35,1.71)* | 1.66 (1.3,2.11)* | 1.17 (0.94,1.47) | 1.21 (1,1.48) | 1.28 (0.98,1.65) | 1.24 (1.04,1.49)* |
|  | Adjusted | 1.48(1.3,1.69)* | 1.56 (1.22,1.99)* | 1.11 (0.88,1.4) | 1.15 (0.94,1.41) | 1.19 (0.92,1.55) | 1.17 (0.97,1.41) |
|  | Adjusted complete cases | 1.35(1.15,1.59)* | 1.42 (1.02,1.97)* | 1.06 (0.77,1.44) | 1.11 (0.85,1.46) | 1.19 (0.85,1.65) | 1.07 (0.83,1.38) |
|  | Gender interaction | **0.009** | 0.971 | 0.276 | 0.491 | 0.28 | 0.193 |
|  | Boys | 1.25(1.05,1.5)* | 1.56 (1,2.44) | 0.96 (0.68,1.36) | 1.25 (0.95,1.65) | 1.02 (0.68,1.54) | 1.01 (0.77,1.34) |
|  | Girls | 1.78(1.48,2.15)* | 1.55 (1.15,2.09)* | 1.23 (0.88,1.72) | 1.02 (0.74,1.42) | 1.31 (0.92,1.87) | 1.24 (0.96,1.61) |
| **Violence between parents** | Basic | 1.59(1.4,1.81)* | 1.07 (0.78,1.47) | 1.21 (0.93,1.58) | 1.71 (1.34,2.2)* | 1.15 (0.84,1.58) | 1.56 (1.27,1.92)* |
|  | Adjusted | 1.14(0.98,1.33) | 0.86 (0.62,1.22) | 1.1 (0.83,1.46) | 1.54 (1.18,1.99)* | 1.02 (0.73,1.42) | 1.36 (1.09,1.69)* |
|  | Adjusted complete cases | 0.96(0.79,1.15) | 1.03 (0.67,1.54) | 0.98 (0.65,1.44) | 1.64 (1.2,2.23)* | 1.04 (0.67,1.58) | 1.47 (1.09,1.97)* |
|  | Gender interaction | 0.536 | 0.371 | 0.761 | 0.731 | 0.883 | 0.963 |
|  | Boys | 1.09(0.89,1.33) | 0.62 (0.29,1.32) | 1.05 (0.69,1.62) | 1.7 (1.16,2.49)* | 1.1 (0.64,1.88) | 1.36 (0.96,1.92) |
|  | Girls | 1.2(0.97,1.5) | 0.97 (0.66,1.43) | 1.14 (0.77,1.7) | 1.41 (0.99,2) | 0.98 (0.63,1.53) | 1.31 (0.97,1.78) |
| **Substance use household** | Basic | 1.72(1.49,2)* | 1.31 (0.89,1.93) | 1.68 (1.23,2.29)* | 2.55 (1.99,3.28)* | 0.95 (0.61,1.47) | 1.82 (1.4,2.37)* |
|  | Adjusted | 1.07(0.9,1.28) | 0.99 (0.64,1.53) | 1.53 (1.09,2.17)* | 2.18 (1.67,2.86)* | 0.76 (0.48,1.22) | 1.53 (1.15,2.05)* |
|  | Adjusted complete cases | 0.96(0.76,1.2) | 0.99 (0.57,1.63) | 1.65 (1.01,2.59)* | 2.16 (1.47,3.14)* | 0.56 (0.26,1.08) | 1.45 (0.97,2.12) |
|  | Gender interaction | 0.733 | 1 | 0.702 | **0.037** | 0.243 | 0.262 |
|  | Boys | 1.03(0.8,1.33) | 0.88 (0.38,2.04) | 1.58 (0.94,2.65) | 1.69 (1.1,2.61)* | 1.12 (0.59,2.12) | 1.41 (0.89,2.23) |
|  | Girls | 1.12(0.85,1.48) | 1.04 (0.61,1.75) | 1.51 (0.94,2.44) | 2.8 (1.9,4.11)* | 0.53 (0.26,1.09) | 1.71 (1.18,2.47)* |
| **Parental mental health problems or suicide** | Basic | 1.57(1.42,1.74)* | 1.69 (1.33,2.14)* | 1.15 (0.93,1.43) | 1.49 (1.24,1.8)* | 1.19 (0.93,1.51) | 1.42 (1.2,1.68)* |
|  | Adjusted | 1.2(1.06,1.36)* | 1.45 (1.11,1.9)* | 1 (0.79,1.27) | 1.39 (1.13,1.72)* | 1.09 (0.82,1.44) | 1.28 (1.05,1.55)* |
|  | Adjusted complete cases | 1.13(0.98,1.3) | 1.26 (0.91,1.76) | 0.8 (0.58,1.09) | 1.42 (1.09,1.85)* | 0.97 (0.69,1.35) | 1.19 (0.93,1.52) |
|  | Gender interaction | 0.838 | 0.293 | 0.41 | 0.131 | 0.21 | 0.836 |
|  | Boys | 1.2(1.01,1.43)* | 1.07 (0.64,1.79) | 1.13 (0.79,1.62) | 1.23 (0.91,1.67) | 0.96 (0.61,1.5) | 1.41 (1.05,1.89)* |
|  | Girls | 1.2(1.01,1.42)* | 1.62 (1.18,2.22)* | 0.9 (0.64,1.27) | 1.57 (1.16,2.11)* | 1.16 (0.8,1.66) | 1.17 (0.91,1.51) |
| **Parent convicted offence** | Basic | 1.79(1.48,2.15)* | 1.44 (0.96,2.14) | 1.37 (0.94,1.99) | 1.57 (1.14,2.16)* | 1.03 (0.63,1.68) | 1.47 (1.07,2.01)* |
|  | Adjusted | 1.27(1.04,1.56)* | 1.2 (0.79,1.83) | 1.33 (0.9,1.95) | 1.39 (0.98,1.96) | 0.85 (0.51,1.4) | 1.29 (0.93,1.8) |
|  | Adjusted complete cases | 1.24(0.96,1.59) | 1.29 (0.73,2.15) | 1.74 (1.07,2.74)* | 1.51 (0.97,2.27) | 0.79 (0.39,1.44) | 1.2 (0.78,1.79) |
|  | Gender interaction | 0.62 | 1 | 0.844 | 0.827 | 0.125 | 0.697 |
|  | Boys | 1.21(0.9,1.61) | 1.16 (0.51,2.63) | 1.32 (0.73,2.4) | 1.46 (0.9,2.38) | 0.48 (0.17,1.37) | 1.26 (0.76,2.08) |
|  | Girls | 1.35(1.02,1.78)* | 1.22 (0.74,2.01) | 1.34 (0.79,2.27) | 1.36 (0.84,2.22) | 1.12 (0.62,2.03) | 1.36 (0.88,2.1) |
| **Parental separation** | Basic | 1.83(1.64,2.05)* | 1.54 (1.19,1.99)* | 1.27 (0.99,1.63) | 1.59 (1.29,1.96)* | 1.21 (0.92,1.58) | 1.85 (1.53,2.23)* |
|  | Adjusted | 1.23(1.08,1.4)* | 1.26 (0.95,1.68) | 1.18 (0.89,1.56) | 1.4 (1.11,1.76)* | 0.97 (0.72,1.31) | 1.58 (1.29,1.95)* |
|  | Adjusted complete cases | 1.2(1.02,1.41)* | 1.12 (0.78,1.61) | 0.93 (0.63,1.32) | 1.49 (1.11,1.99)* | 1 (0.68,1.44) | 1.56 (1.19,2.03)* |
|  | Gender interaction | 0.368 | 1 | 0.741 | 0.808 | 0.109 | 0.926 |
|  | Boys | 1.32(1.11,1.57)* | 1.16 (0.65,2.08) | 1.31 (0.87,1.96) | 1.56 (1.11,2.18)* | 0.72 (0.42,1.24) | 1.68 (1.2,2.36)* |
|  | Girls | 1.16(0.96,1.41) | 1.31 (0.93,1.83) | 1.1 (0.76,1.59) | 1.29 (0.91,1.84) | 1.14 (0.78,1.67) | 1.56 (1.18,2.06)* |

*Supplemental Table 6 P-values for the interaction between each ACE and parental social class.*

| **Adversity** | **Analysis 1: Education** | **Analysis 2: Health** | | | | |
| --- | --- | --- | --- | --- | --- | --- |
|  | < Five GCSEs | Depression | Harmful alcohol use | Illicit drug use | Obesity | Smoking |
| Categorical classic ACEs 1 | 0.85 | 0.60 | 0.51 | 0.11 | 0.61 | 0.35 |
| 2-3 | 0.33 | 0.99 | 0.55 | 0.09 | 0.74 | 0.37 |
| 4+ | 0.44 | 0.82 | 0.69 | 0.46 | 0.69 | 0.22 |
| Physical abuse | 0.85 | 0.61 | 0.98 | 0.88 | 0.25 | 0.68 |
| Sexual abuse | 0.68 | 0.54 | 0.14 | 0.53 | 0.69 | 0.47 |
| Emotional abuse | 0.69 | 0.93 | 0.83 | 0.99 | 0.76 | 0.56 |
| Emotional neglect | 0.95 | 0.67 | 0.81 | 0.07 | 0.90 | 0.34 |
| Bullying | 0.26 | 0.81 | 0.54 | 0.42 | 0.35 | 0.49 |
| Violence between parents | 0.99 | 0.99 | 0.89 | 0.46 | 0.68 | 0.28 |
| Substance household | 0.74 | 0.68 | 0.53 | 0.69 | 0.77 | 0.32 |
| Mental health problems or suicide | 0.68 | 0.92 | 0.85 | 0.73 | 0.77 | 0.37 |
| Parent convicted offence | 0.84 | 0.90 | 0.37 | 0.91 | 0.87 | 0.58 |
| Parental separation | 0.21 | 0.85 | 0.67 | 0.46 | 0.96 | 0.81 |

*Supplemental Table 7 P-values for the interaction between each ACE and maternal education.*

| **Adversity** | **Analysis 1: education** | | **Analysis 2: health** | | | | |
| --- | --- | --- | --- | --- | --- | --- | --- |
|  | Less than five GCSEs | Depression | | Harmful alcohol use | Illicit drug use | Obesity | Smoking |
| Categorical classic ACEs 1 | 0.24 | 0.38 | | 0.81 | 0.92 | 0.35 | 0.09 |
| 2-3 | 0.06 | 0.24 | | 0.31 | 0.75 | 0.18 | 0.10 |
| 4+ | 0.60 | 0.48 | | 0.39 | 0.80 | 0.40 | 0.10 |
| Physical abuse | 0.29 | 0.52 | | 0.91 | 0.57 | 0.55 | 0.98 |
| Sexual abuse | 0.36 | 0.52 | | 0.40 | 0.66 | 0.73 | 0.38 |
| Emotional abuse | 0.05 | 0.73 | | 0.50 | 0.38 | 0.13 | 0.15 |
| Emotional neglect | 0.94 | 0.24 | | 0.25 | 0.51 | 0.98 | 0.65 |
| Bullying | 0.72 | 0.52 | | 0.73 | 0.20 | 0.81 | 0.45 |
| Violence between parents | 0.30 | 0.72 | | 0.28 | 0.81 | 0.85 | 0.59 |
| Substance household | 0.31 | 0.96 | | 0.76 | 0.96 | 0.87 | 0.44 |
| Mental health problems or suicide | 0.46 | 0.53 | | 0.84 | 0.45 | 0.92 | 0.14 |
| Parent convicted offence | 0.10 | 0.95 | | 0.57 | 0.30 | 0.73 | 0.12 |
| Parental separation | 0.08 | 0.74 | | 0.96 | 0.73 | 0.68 | 0.46 |

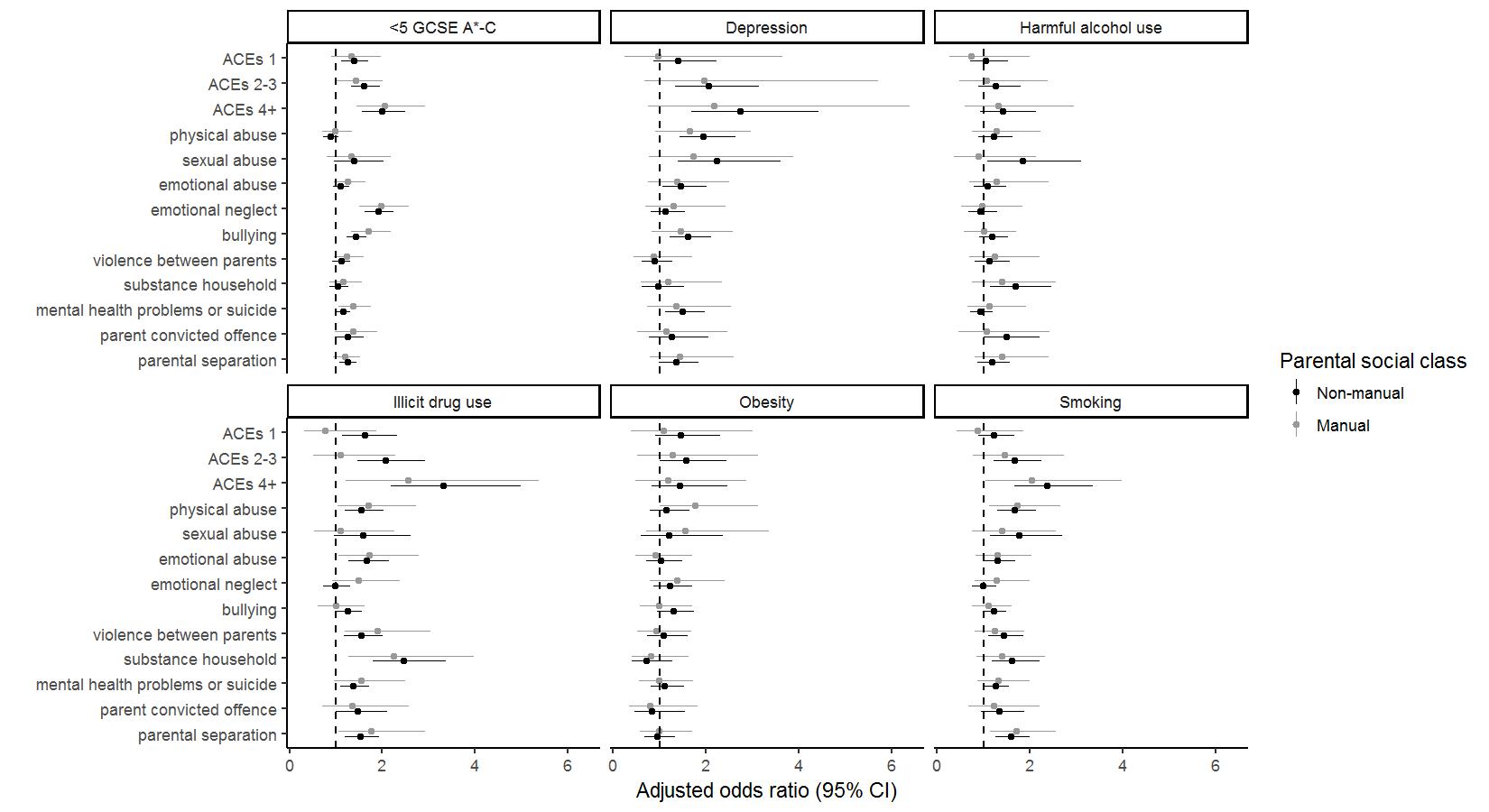

Supplementary Figure 1: Associations between ACE and educational attainment (less than 5 GCSEs), health or health-related behaviours (Depression, Harmful alcohol use, Illicit drug use, Obesity, Smoking), stratified by parental social class.

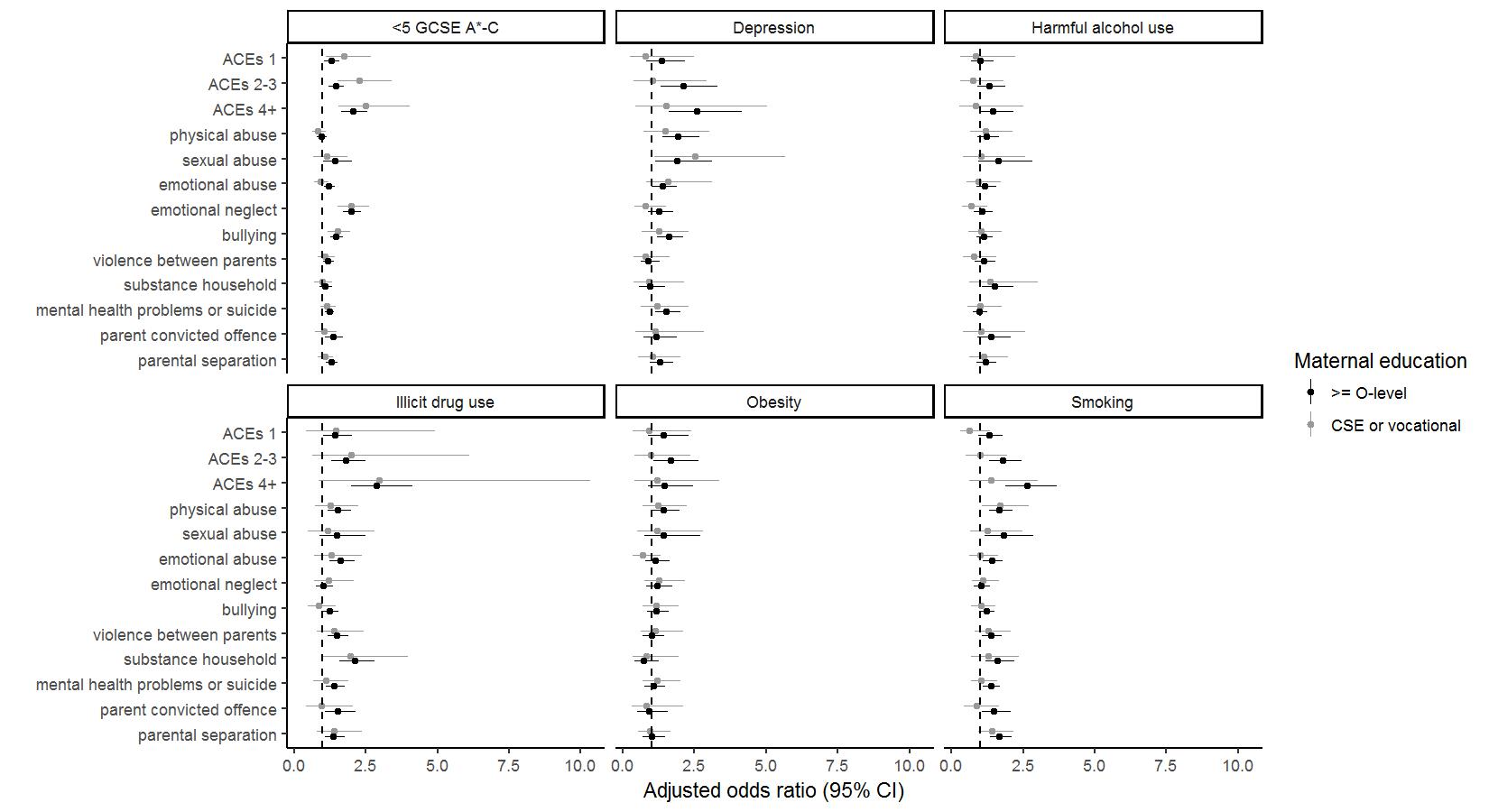

Supplementary Figure 2: Associations between ACE and educational attainment (less than 5 GCSEs), health or health-related behaviours (Depression, Harmful alcohol use, Illicit drug use, Obesity, Smoking), stratified by maternal education.
